# TRAP1 *S*-nitrosylation as a model of population-shift mechanism to study the effects of nitric oxide on redox-sensitive oncoproteins

**DOI:** 10.1101/2022.12.11.519943

**Authors:** Elena Papaleo, Matteo Tiberti, Matteo Arnaudi, Chiara Pecorari, Fiorella Faienza, Lisa Cantwell, Kristine Degn, Francesca Pacello, Andrea Battistoni, Matteo Lambrughi, Giuseppe Filomeni

**Affiliations:** Cancer Structural Biology, Danish Cancer Society Research Center, 2100, Copenhagen, Denmark; Cancer Systems Biology, Section for Bioinformatics, Department of Health and Technology, Technical University of Denmark, 2800, Lyngby, Denmark; Redox Biology, Danish Cancer Society Research Center, 2100, Copenhagen, Denmark; Department of Biology, University of Rome Tor Vergata, 00133, Rome, Italy; Center for Healthy Aging, Copenhagen University, 2200, Copenhagen, Denmark

**Keywords:** redox switch, TRAP1, disulfide, cysteine, molecular simulations, redox signaling, bioinformatics

## Abstract

*S*-nitrosylation is a post-translational modification in which nitric oxide (NO) binds to the thiol group of cysteine, generating an *S*-nitrosothiol (SNO) adduct. *S*-nitrosylation has different physiological roles, and its alteration has also been linked to a growing list of pathologies, including cancer. SNO can affect the function and stability of different proteins, such as the mitochondrial chaperone TRAP1. Interestingly, the SNO site (C501) of TRAP1 is in the proximity of another cysteine (C527). This feature suggests that the *S*-nitrosylated C501 could engage in a disulfide bridge with C527 in TRAP1, resembling the well-known ability of *S*-nitrosylated cysteines to resolve in disulfide bridge with vicinal cysteines. We used enhanced sampling simulations and in-vitro biochemical assays to address the structural mechanisms induced by TRAP1 *S-*nitrosylation. We showed that the SNO site induces conformational changes in the proximal cysteine and favors conformations suitable for disulfide-bridge formation. We explored 4172 known *S*-nitrosylated proteins using high-throughput structural analyses. Furthermore, we carried out coarse-grain simulations of 44 proteins to account for protein dynamics in the analyses. This resulted in the identification of up to 1248 examples of proximal cysteines which could sense the redox state of the SNO site, opening new perspectives on the biological effects of redox switches. In addition, we devised two bioinformatic workflows (https://github.com/ELELAB/SNO_investigation_pipelines) to identify proximal or vicinal cysteines for a SNO site with accompanying structural annotations. Finally, we analyzed mutations in tumor suppressor or oncogenes in connection with the conformational switch induced by *S*-nitrosylation. We classified the variants as neutral, stabilizing, or destabilizing with respect to the propensity to be *S*-nitrosylated and to undergo the population-shift mechanism. The methods applied here provide a comprehensive toolkit for future high-throughput studies of new protein candidates, variant classification, and a rich data source for the research community in the NO field.

## Introduction

Nitric oxide (NO) can exert its physiological effects through post-translational modification (PTM) of protein thiol groups. This modification is known as *S-*nitrosylation (SNO), which occurs when an NO moiety reacts with the thiol group of cysteine.^1^

*S-*nitrosylation is a PTM regulating protein function, it is present in all kingdoms of life^1,2^, and it is often referred to as the “prototypic” redox-based signaling mechanism^1^. Furthermore, *S-* nitrosylation is a crucial mediator of NO signaling. This PTM is involved in various physiological processes, as it regulates metabolic enzymes, oxidoreductases, proteases, phosphatases, protein kinases, respiratory complexes, receptors, ion channels, transporters, cytoskeletal components, and transcription factors^3^. More than 4000 experimentally characterized S-nitrosylated cysteines have been reported in a database dedicated to PTMs (dbPTM)^4^.

NO overproduction, as an effect of increased expression of the inducible form of nitric oxide synthase (iNOS), has been associated with aggressive tumor phenotypes and poor outcomes in different cancer types ^5^. If this depends, at least in part, on *S-*nitrosylation is still argued. However, aberrant *S-*nitrosylation has been reported in aggressive breast cancer^6^ and hepatocellular carcinoma ^7,8^. These alterations are due to the downregulation or deletion of the best-characterized denitrosylase, *S-*nitrosoglutathione reductase (GSNOR). GSNOR removes the NO group from *S-*nitrosoglutathione (GSNO), which is in equilibrium with *S-* nitrosylated proteins., thus deactivating the effects of *S-*nitrosylation on proteins.^9^ In cellular models of hepatocellular carcinoma, GSNOR downregulation, or loss, induces the *S-* nitrosylation of the mitochondrial chaperone TRAP1 ^10,11^ at Cys501 (numbering in the human variant). *S-*nitrosylation of TRAP1 results in its accelerated degradation via the proteasome.^7^ Besides acting as a regulatory PTM, *S-*nitrosylation can make a cysteine susceptible to the nucleophilic attack of the sulfhydryl group of a neighboring reduced cysteine, leading to the formation of a disulfide bond and the release of the NO moiety, as nitroxyl anion (NO^-^)^1,12^. Disulfide bond (or bridge) formation, in turn, can increase or decrease structural stability, depending on the context ^13^.

To our knowledge, the formation of a disulfide bridge after *S-*nitrosylation has only been observed between vicinal cysteines, i.e., two cysteines that are both spatially adjacent and close in sequence with a CXXC sequence motif. A well-characterized example of this is thioredoxin ^14–16^.

However, after *S-*nitrosylation, a disulfide bridge might also be generated between an *S-* nitrosylated cysteine and a proximal cysteine, i.e., cysteines within the same protein are close in the three-dimensional (3D) structure, but not in sequence. An example of this class is the aforementioned mitochondrial chaperone, TRAP1, an oncoprotein, in which the SNO site (Cys501) has been experimentally characterized with Cys527 in its proximity (**Figure 1A**). In this context, molecular modeling and simulations are promising tools for investigating the structural mechanisms induced by *S-*nitrosylation ^3^.

**Figure 1.**
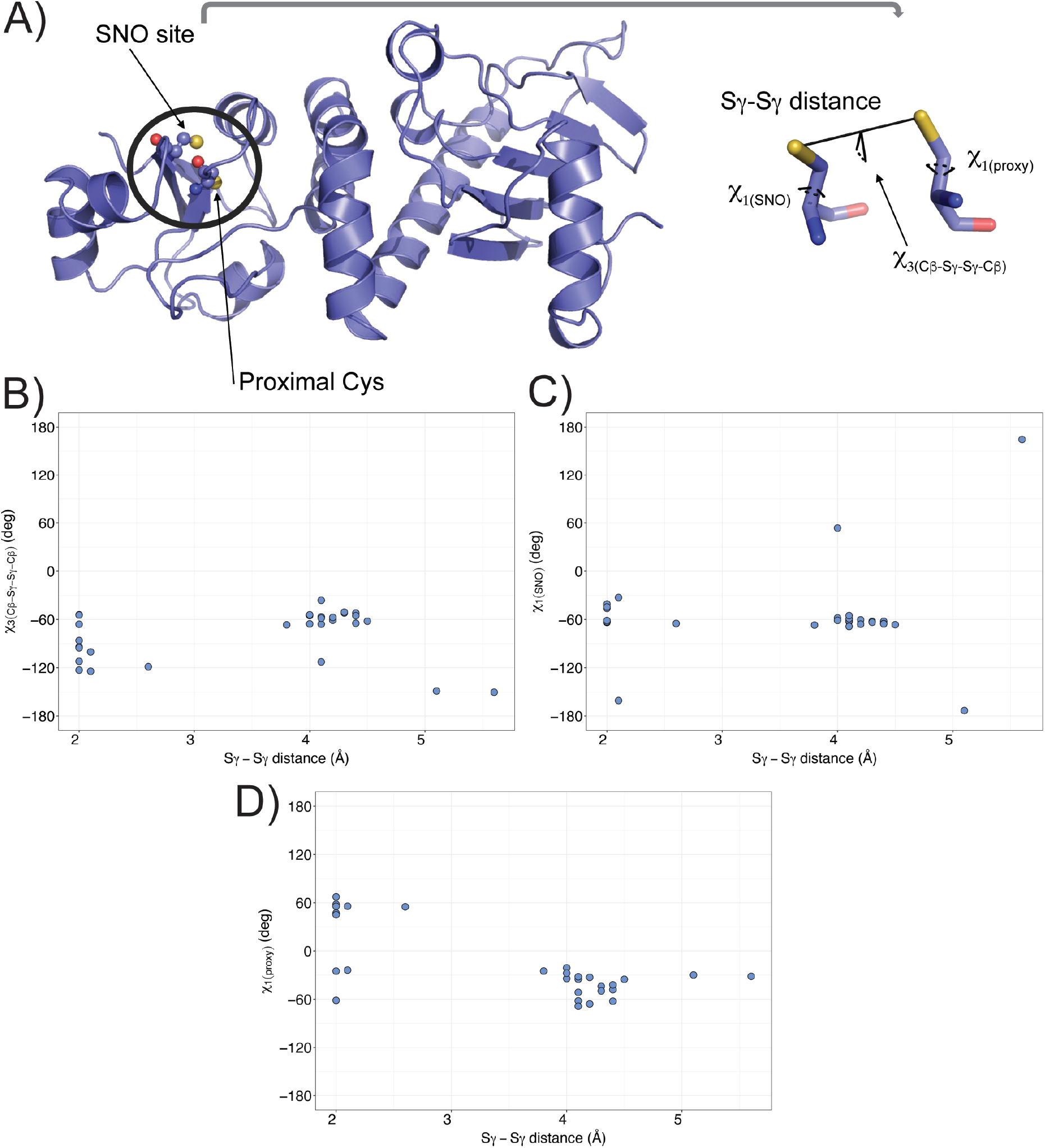
Conformational heterogeneity of the *S-*nitrosylated cysteine and its proximal cysteine in the mitochondrial chaperone TRAP1. A) In the left panel, the cartoon shows the X-ray structure of the middle domain of *Danio rerio* TRAP1 (PDB ID entry 4IPE, residues 311-567). The spheres indicate the atoms of the *S-*nitrosylated (SNO) site (C516) and the proximal cysteine (C542). In the right panel, the stick representation shows a conformation of the SNO site and the proximal cysteine. We evaluated the conformational state and orientation of each cysteine, measuring i) the χ1 dihedral angle of the SNO cysteine (χ_1(SNO)_), ii) the χ1 dihedral angle of the proximal cysteine (χ_1(proxy)_), iii) the distance between the two sulfur atoms of the two cysteines (Sγ-Sγ distance), and iv) the Cβ-Sγ-Sγ-Cβ dihedral angle. B-D) We retrieved from the Protein Data Bank (PDB) the structures available for *Homo Sapiens* and *Danio rerio* TRAP1. We evaluated the conformational state of the *S*-nitrosylated cysteine (C501 and C516, respectively) and its proximal cysteine (C527 and C542, respectively). The scatter plots show the calculated values for each structure of the Sγ-Sγ distance against B) Cβ-Sγ-Sγ-Cβ, C) χ_1(SNO)_, and D) χ_1(proxy)_ dihedral angles. We observed heterogeneity in the conformation and orientation of the two cysteines, showing Sγ-Sγ distances in the range of 2-5.6 Å and diversity in Cβ-Sγ-Sγ-Cβ (in the range between −150.5 and −36.2°), χ_1(SNO)_ (in the range between −173.3 and 164.5°), and χ_1(proxy)_ (in the range between −68.7 and 55.0°) dihedral angles.

In light of the above observations, we here applied molecular simulations and bioinformatic analyses to investigate whether the *S*-nitrosylation of TRAP1 could induce local structural changes in an SNO site or its proximal cysteine. Our work shows evidence of a population-shift mechanism in which the proximal site acts as a redox switch that senses the state of the SNO site. We then analyzed the *S*-nitrosylated proteome using a bioinformatic workflow designed for this purpose (SNOfinder) and identified 631 proteins from different organisms that can undergo similar mechanisms. Of these, 95 SNO sites were found in human proteins, among which we identified five cancer-driver proteins of interest (TP53, GRIN2A, CBFB, CALR, and EGFR). Thus, we further investigated these proteins to understand if any known cancer mutation, which falls in the proximity of the SNO site, or the proximal cysteines, could alter the population shift induced by SNO.

Our work sheds new light on the mechanisms induced by SNO modifications of proteins, including multiple cysteines. We also provide a toolkit that can be applied to characterize newly reported events of *S*-nitrosylation for their capability to favor disulfide bridges through a population-shift mechanism and predict the effects of disease-related mutations in the surroundings of the SNO site.

## Results

### Conformational heterogeneity of the SNO site and its proximal cysteine in TRAP1 X-ray structures

At first, we retrieved the available structures for the human and *Danio rerio* orthologs of TRAP1 from the Protein Data Bank (PDB). We included TRAP1 structures from *Danio rerio* as we used this ortholog as a model system in our previous study ^17^. We retained only the structures, which included the coordinates for the *S-*nitrosylated cysteine (C501 and C516 in human and *Danio rerio*, respectively) and its proximal cysteine (C527 and C542, respectively). We also verified that the PDBREDO refinement protocol ^18^ would not alter the conformation of the SNO site and proximal cysteine before starting the analyses. We then evaluated the conformational state of each cysteine and reciprocal orientation, measuring the properties reported in **Table S1** and **Figure 1 B-D**.

We noticed that, in some X-ray structures, the electron density in the region around the proximal cysteine suggested two alternative conformations, one at a distance of approximately 2 Å of the SNO site and the other 4 Å afar, where the two cysteines point in different directions **(Table S1)**. Overall, in the set of analyzed structures, the two cysteines were at distances in the 2-5.6 Å range, suggesting that different conformational states can be at play in this region. We also identified a similar heterogeneity for the Cβ-Sγ-Sγ-Cβ dihedral angle and the χ_1_ of the two cysteines, especially for the proximal cysteine residues (**Table S1, Figure 1B-D**). Some of the observed conformations satisfied known criteria for disulfide bridge formation, i.e., sulfur-to-sulfur distances of 2.5 Å or lower ^19^ and Cβ-Sγ-Sγ-Cβ dihedral angles around ±90 degrees ^20^.

### Molecular dynamics investigation of the SNO site and its proximal cysteine of TRAP1 in their reduced and S-nitrosylated forms

We collected different simulations using enhanced sampling for the middle domain of the *Danio rerio* TRAP1_311-567_ variant, as reported in **Table S2**. To understand if the *S*-nitrosylation of C516 could induce conformational changes on the proximal cysteine C542, we estimated the free energy profile associated with different collective variables depicted in **Figure 1A**. The collective variables have been designed to capture the conformation of the proximal cysteine and its surroundings (see Materials and Methods). In particular, we used the χ_1_ of the two cysteine residues (i.e., χ_1(SNO)_ and χ_1(proxy)_), the distance between the two sulfur groups of the two cysteines (Sγ-Sγ distance), and the Cβ-Sγ-Sγ-Cβ dihedral angle. Similarly, we also included in the analyses an unbiased one-μs molecular dynamics simulation of the oxidized form of TRAP1 carrying a disulfide bridge between the two cysteines, to be used as a reference for the measurements that we use as collective variables.

### Different molecular dynamics force fields agree on the structural features of the SNO and proximal site

Molecular dynamics physical models (i.e., force fields) are known to be very sensitive in terms of their capability to describe residue rotameric states and local conformations of protein residues. However, a consensus on the dynamics described by different force fields for the same protein can often be difficult to achieve ^21–23^. Thus, before comparing *S-*nitrosylated and reduced variants of TRAP1 – for which parameters are available only for a small number of force fields – we scrutinized eight different force fields (**Table S2**). We focused on the capability of the different force field to describe the conformations of the reduced cysteine. To this end, we applied an enhanced sampling method based on metadynamics and the same collective variables described in the previous section.

Most simulations showed that, when fully reduced, both cysteines populate the three possible rotameric states associated with their χ_1_ (trans(t), minus(m), and plus(p)), with the p rotamer as the less favorable one (**Figure 2A-B**). However, in the GROMOS54a7 simulations, the proximal cysteine seems to equally populate all the dihedral states (**Figure 2B**). We then observed that GROMOS54a7 also stood out considering the distance between the two cysteine residues **(Figure 2C**) as the only simulation with a free energy minimum at distances greater than 7 Å. The mono-dimensional free energy profiles for the Cβ-Sγ-Sγ-Cβ dihedral were more variable for the different force fields but still able to captured the same main minima (**Figure S1**). We observed that GROMOS54a7 was also an outlier in this case, with a less pronounced energy barrier among the states.

**Figure 2.**
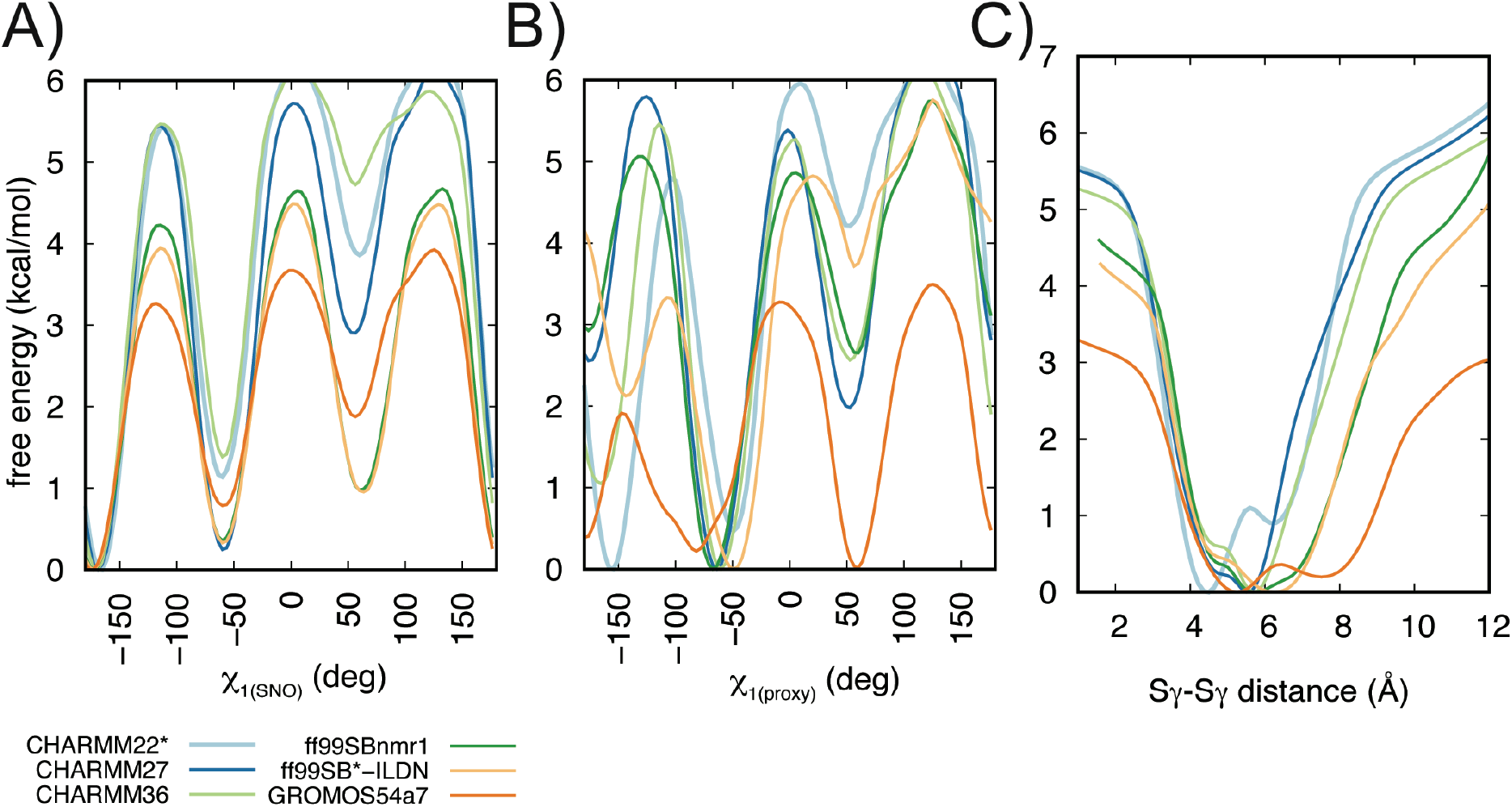
Different MD force fields agree on the conformational preferences of the reduced-form of the two cysteines of TRAP1_311-567_. The panels show the mono-dimensional free energy profiles associated with the collective variables A) χ_1(SNO)_, B) χ_1(proxy)_ and C) Sγ-Sγ distance for the metadynamics simulations. We performed metadynamics using CHARMM22* (light blue), CHARMM27 (dark blue), CHARMM36 (light green), ff99SBnmr1 (dark green), ff99SB*-ILDN (light orange), GROMOS54a7 (dark orange) force fields. Overall, the MD force fields agree that the SNO site and the proximal cysteine could populate three possible rotameric states for their χ_1_ dihedrals, with the plus state as the less favorable one. Furthermore, they agree in describing the distances between the two cysteines with minima in the free energy landscape at a distance of 4-6.5 Å. We observed that GROMOS54a7 is the only exception showing a less stable behavior, with χ_1(proxy)_ equally populating all the dihedral states and the Sγ-Sγ distance sampling minima higher than 7 Å.

### S-nitrosylation induces a population shift mechanism on the proximal cysteine and favors conformations enabling the formation of a disulfide bridge

Using the same enhanced sampling approach applied above, we simulated the *S*-nitrosylated variant at the C516 site. At the time of this study, we identified parameters for *S*-nitrosylated cysteines that could be used in combination with ff99SB*-ILDN ^24,25^ or GROMOS54a7 ^26^. As the simulation of the reduiced form of TRAP1 with GROMOS54a7 (**Figure 2A-C, Figure S1**) showed a less stable behavior, we relied on the results from ff99SB*-ILDN (**Figure 3**).

**Figure 3.**
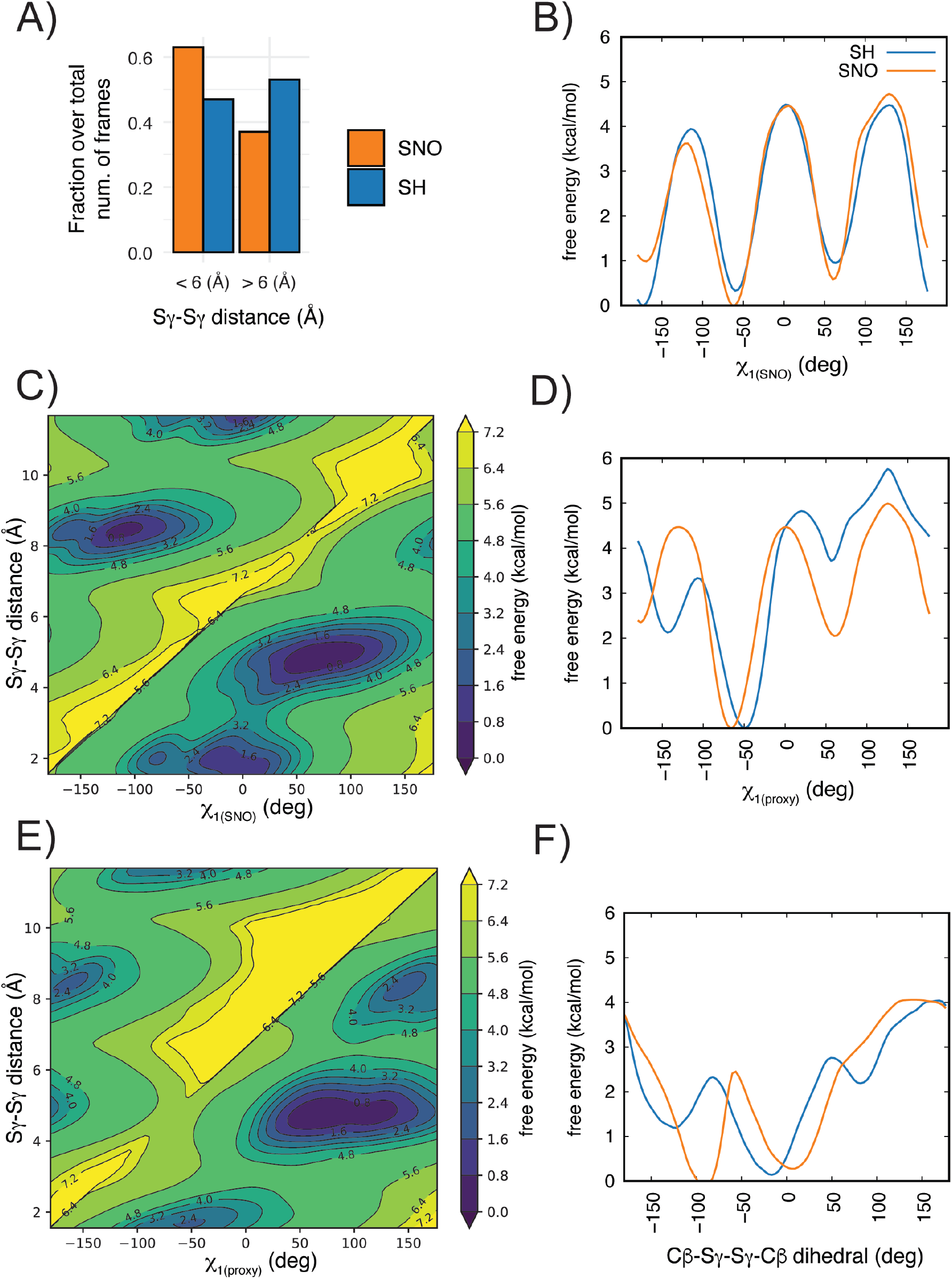
*S*-nitrosylation of C516 of TRAP1 favors conformations more suited to the disulfide-bridge formation with the proximal cysteine. We performed metadynamics of the reduced (SH, blue) and *S*-nitrosylated (SNO, orange) variants at the C516 site of *Danio Rerio* TRAP1_311-567_ using ff99SB*-ILDN force field. A) The bar plot shows the fraction over the total number of frames of the metadynamics trajectories with Sγ-Sγ distance values lower/higher than 6 Å. We observed a higher propensity towards lower distances between the two cysteines upon *S-*nitrosylation of C516. B-F) Mono- and bi-dimensional free energy surfaces associated with the collective variables for the metadynamics simulations: B) χ_1(SNO)_, C) χ_1(SNO)_ and Sγ-Sγ distance, D) χ_1(proxy)_, E) χ_1(proxy)_ and Sγ-Sγ distance, F) Cβ-Sγ-Sγ-Cβ. We observed that the *S*-nitrosylation of C516 disfavors the trans states of χ_1(SNO)_ of around 1 kcal/mol (panel A). The *S*-nitrosylation of C516 disfavors the trans states of χ_1(proxy)_ of more than 2 kcal/mol and makes more favorable the plus states of χ_1(proxy)_ (panel D). The plus and minus states of χ_1(SNO)_ and χ_1(proxy)_ correlate with Sγ-Sγ distances lower than 6 Å between the two cysteines (panels C and E). *S*-nitrosylation favors a conformation for the Cβ-Sγ-Sγ-Cβ dihedral angle around −90 degrees which is often observed in disulfide bonds (panel F).

As a reference, we evaluated the same collective variables used in metadynamics in a reference simulation of a TRAP1 variant in which the disulfide bridge is formed (**Figure S2**). In the reference simulations, the *S*-nitrosylated and the proximal cysteine featured a m and p rotamer for their side chain, respectively. We found the Cβ-Sγ-Sγ-Cβ dihedral to be frequently found at a value close to −90 degrees in our simulation, which is often observed in disulfide bridges, and the distance between the two sulfur atoms is 2.03 Å.

When comparing the two simulations with reduced (SH) or *S-*nitrosylated C516, we observed that lower distances between the two cysteines are preferred upon *S-*nitrosylation (**Figure 3A**). When we measured the distance between the NO group and the proximal cysteine, we observed even lower distances (**OSF repository**, https://osf.io/sfrkc/).

This preference is not explained entirely by the changes in the SNO site. Indeed, the *S*-nitrosylation did not have a marked effect on the conformation of the SNO site apart from disfavoring the t conformer of the χ_1_ of 1 kcal/mol (**Figure 3B**). Interestingly, the p and m rotamers for the χ_1_ of the *S*-nitrosylated C516 are also the most populated when the two cysteines are close by (**Figure 3C**). The main effect imparted by *S-*nitrosylation is on the proximal cysteine where the t rotamer for χ_1_ is disfavoured of more than 2 kcal/mol, and the p state becomes more favorable than in the reduced variant of the protein of approximately the same amount. Nevertheless, The proximal cysteine could still explore the m χ_1_ rotamer upon *S*-nitrosylation (**Figure 3D**). The p and m states seem to correlate with distances lower than 6 Å between the two cysteines, whereas the t rotamer for χ_1_ of C542 is found mainly at distances higher than 8 Å (**Figure 3E**). Moreover, *S-*nitrosylation favors a geometry for the Cβ-Sγ-Sγ-Cβ dihedral angle around −90 degrees (**Figure 3F**). This agrees with values for this angle that, for canonical disulfide bonds, is ±90 degrees ^27^.

### Exposure to nitric oxide induces the formation of a disulfide bridge between C501 and C527 in human recombinant TRAP1

We previously demonstrated that NO fluxes induce recombinant human TRAP1 *S-* nitrosylation at C501. These results were also confirmed in a cell system in which the loss of the GSNOR denitrosylate increases *S-*nitrosylation.^7^ Based on this evidence and the above-reported simulations, we wanted to experimentally verify if C501 and C527 could form a disulfide bond upon NO exposure. Other than C501 and C527, TRAP1 has two further cysteines, C261 and C573, which are exposed to the solvent and can potentially react with electrophiles. Therefore, to rule out any contribution of these residues, we generated a C261S-C573R mutant of human TRAP1 (C_2_-TRAP1), in which C501 and C527 were the only cysteines that could react and be engaged in a disulfide bridge. The mutations introduced are predicted as neutral in terms of structural stability using free energy calculations (ΔΔG ∼ 0 kcal/mol), suggesting that the mutated variant C_2_-TRAP1 maintains its structural integrity.

C_2_-TRAP1 was next incubated with an NO donor *S*-nitroso-*N*-acetylpenicillamine (SNAP) or DTT (as control) and the redox modifications induced (*S*-nitrosylation or disulfide bridge) were analyzed by Mal-PEG switch assay. This methodology relies on a sequence of three reactions exemplified as in **Figure 4A**; namely: i) Blocking – usually *via* alkylation – of free (reduced) sulfhydryls; ii) Reduction of the oxidized sulphur atoms (i.e., cysteine residues engaged in SNO or S-S bound upon SNAP treatment), and iii) Alkylation of the newly generated thiols with maleimide conjugated with polyethylene-glycol (Mal-PEG), which confers a different mobility shift to the protein on SDS-PAGE. Results show that SNAP increased the apparent molecular weight of the recombinant C_2_-TRAP1 immunoreactive band. Besides the amount of C_2_-TRAP1 that did not react (approximately the 50% of the total protein loaded on the gel), we were able to detect two further bands at higher molecular weights (**Figure 4B**). Of these, the lowest one (which accounted by the 33% of the total protein), indicates a single Mal-PEG addition, reasonably the SNO modification at C501; whereas the highest one (which accounted by the remaining 17%) represents the amount of TRAP1 bound to two Mal-PEG moieties, likely the S-S variant. To exclude any unwanted technical artifact induced by Mal-PEG, we also mutagenized C501 and C527 and produced a cysteine-free variant of TRAP1 (C_0_-TRAP1). We then applied the same treatment and procedures as above, but did not observe any change in the electrophoretic pattern of C_0_-TRAP1 upon SNAP treatment. Altogether, these results confirmed that *S*-nitrosylation of C501 – previously demonstrated to take place upon NO exposure – can resolve in a disulfide bridge with C527.

**Figure 4.**
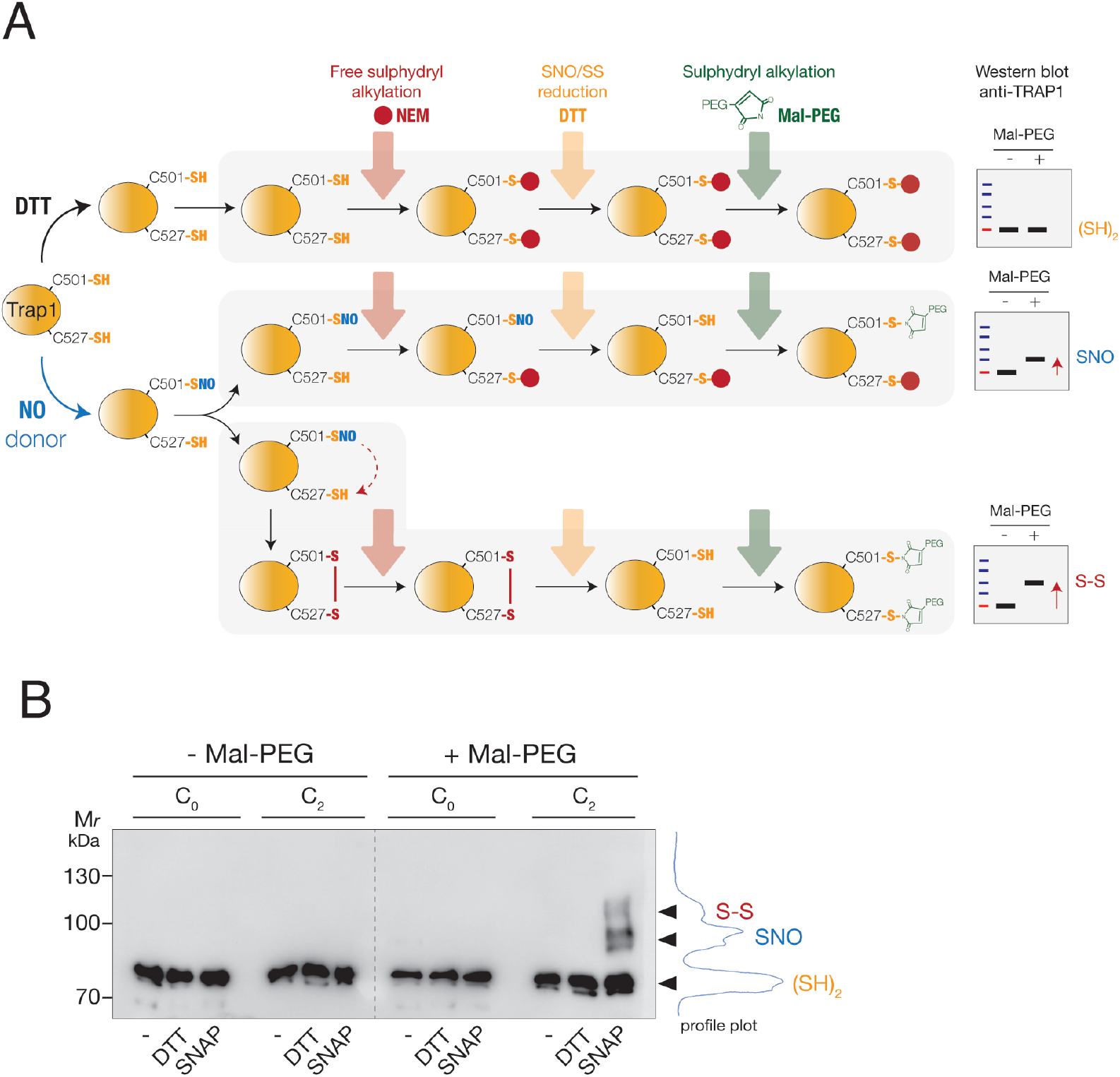
Exposure to NO fluxes results in the formation of a disulfide bridge between C501 and C527 in human recombinant TRAP1. a) Scheme of the protocol applied to detect cysteine redox modifications in human recombinant TRAP1. C_2_-TRAP1 (which possesses only C501 and C527) has been treated in vitro with 500 µM SNAP, or DTT (as control) for 4 h, at RT. Afterward, the excess of SNAP or DTT was washed out, and the protein was sequentially subjected to: i) thiol alkylation with *N*-ethylmaleimide (NEM) (red arrow), followed by ii) reduction with DTT (orange arrow) and iii) alkylation with Mal-PEG (green arrow), which confers an increase of the molecular weight proportional to the number of Mal-PEG bound. Depending on the C501 and C527 oxidation state [i.e., *S*-nitrosylated (SNO), or oxidized to disulfide (S-S)], the sequence of reactions above described generates one or two immunoreactive bands upon Western blot analyses with an anti-TRAP1 antibody, which show apparent molecular weights higher than the fully reduced (SH)_2_ or untreated protein. The same protocol was applied to C_0_ (cysteine-free) mutant of TRAP1 as a negative control. b) Western blot analyses of 1 µg of human recombinant C_0_ and C_2_-TRAP1 mutants treated as described in a). Fully reduced ((SH)_2_), *S*-nitrosylated (SNO) or disulfide-containing (S-S) variant of C_2_-TRAP1 are shown together with the profile plot calculated with Fiji. Note that the molecular weight shift induced by the addition of one or more Mal-PEG moieties is not directly additive to the size of TRAP1. As reported in ^101^, this is due to drag and steric hindrance of branched proteins occurring during the migration through a gel compared to unlabelled linear proteins, which results in a shift which is usually greater than expected.

### A proteome-wide search of other candidates with an S-nitrosylated cysteine and a proximal cysteine

TRAP1 may not be the only example of the mechanism proposed here where the *S*-nitrosylation of one site can induce a conformational change to the proximal cysteine and promote conformations suited to disulfide-bridge formation. We thus devised a bioinformatic pipeline, SNOfinder (**Figure 5A** and Materials and Methods), to find other candidates among the known *S*-nitrosylated proteins reported in dbPTM ^4^. SNOfinder identified 1477 cases of an SNO site with a proximal cysteine according to the distance criteria in 631 proteins. Of these, only 489 sites featured accessibility of the cysteine side chain higher than 10% (**Figure 5B**). The remaining SNO sites were rather buried. Buried PTM sites could become available for their modifying enzymes thanks to conformational dynamics, as observed, for example, for phosphorylation ^28,29^. However, SNOfinder does not include the modelling of protein dynamics. Therefore, we decided to focus on sites that are at least partially solvent accessible and could represent more reliable hits as candidates for modifications by NO. Most sites are also located in regions with confident prediction in the models deposited in the AlphaFold database^30^ (**Figure 5B**). This suggests that they are of high enough quality to be further analyzed with molecular modelling^31^.

**Figure 5.**
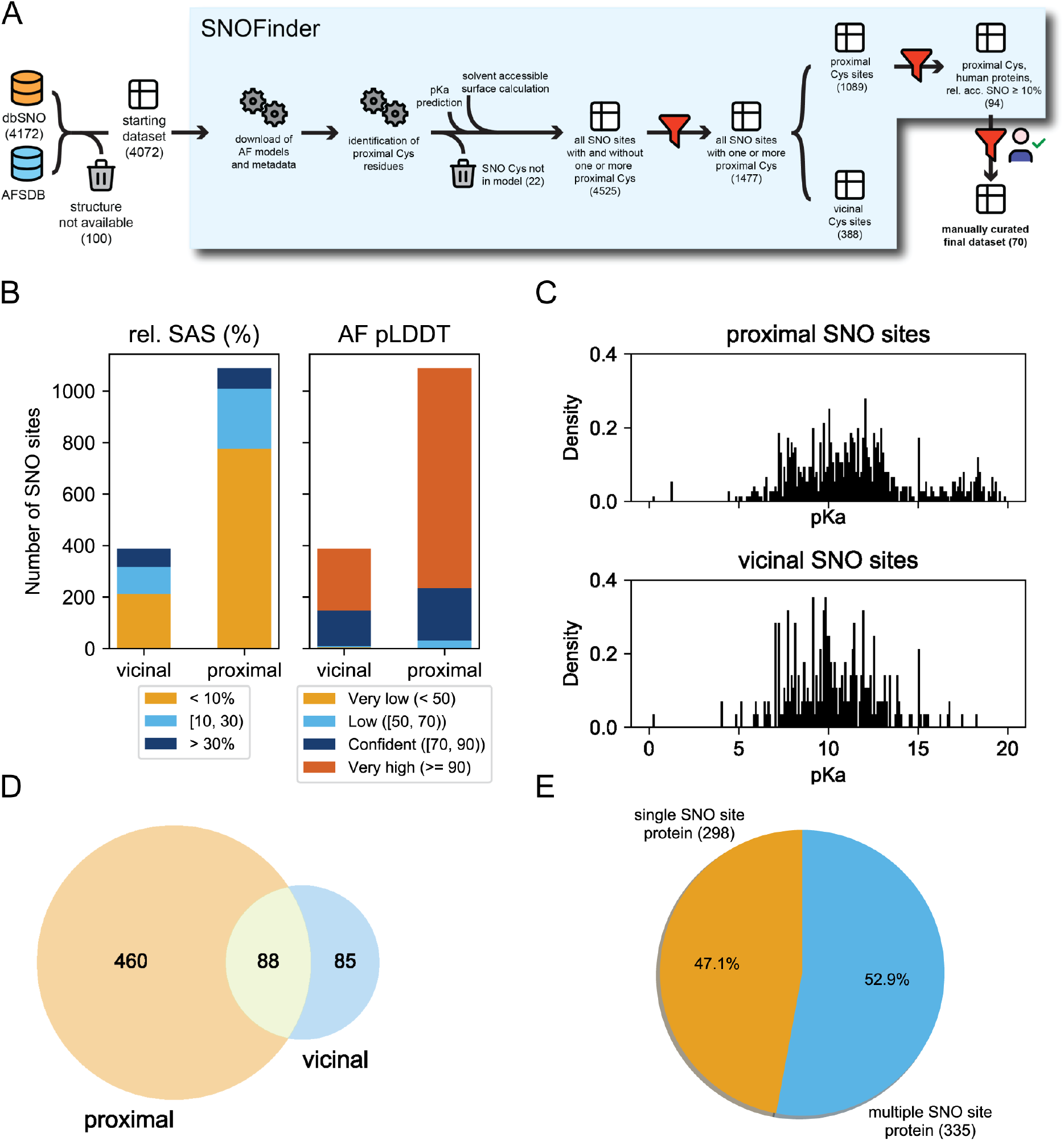
SNOfinder pipeline and analysis of the SNO and proximal cysteine dataset. a) schematic representation of the SNOfinder pipeline. We start from a database of known SNO sites and their proteins (dbPTM). We first identify which of these proteins have available models in the AlphaFold Protein Structure Database (AFPSDB) and only keep those. We then analyze each model to identify proximal cysteines to the SNO sites, as detailed in Methods. For both the SNO site and the proximal cysteine, we annotate predicted pK_a_, AlphaFold pLDDT score, secondary structure assignment, and relative solvent accessible surface (see Methods). We then further filter the obtained dataset by keeping only entries (i.e., SNO sites) with at least one proximal cysteine and split them between vicinal (i.e., the SNO and proximal Cys are at less than eight residues apart in sequence) and proximal (i.e., the two residues are spatially close but not close in sequence). Finally, we generate a dataset in which we keep only human proteins and for which the SNO site has at least 10% solvent-accessible surface area. This dataset was then manually curated (see Methods), obtaining the final dataset. The number in parenthesis under each step denotes the size of the dataset, as a number of SNO sites or SNO sites/Cys pairs B) Classification of the SNO sites as found in the AlphaFold models according to relative solvent accessible surface (left) and AlphaFold pLDDT score (right) C) Obtained distributions of pK_a_ values for SNO site cysteines in the vicinal and proximal SNO datasets. Before plotting, we excluded cases for which the predicted pK_a_ values were unphysical (i.e., pK_a_ = 99.99) D) Venn diagram representation of proteins having SNO sites with distal or proximal cysteine residues E) pie chart of proteins having only one SNO site vs. proteins having multiple SNO sites, among those in which at least one proximal or vicinal cysteine was identified.

As expected, our dataset included TRAP1 C501 and C527 as SNO sites and proximal cysteine, respectively. We then evaluated which sites had *vicinal* cysteines that resembled the thioredoxin case^14,15^ and are close in the sequence (**Figure 5A**, 388 sites overall and 176 partially solvent accessible, **OSF** repository: https://osf.io/52bng/). In addition, we retrieved the ones that resemble the TRAP1 case in which the two cysteines are closed in the folded 3D structure but not in sequence, i.e., *proximal* cysteines (**Figure 5A**, 1089 overall and 313 sites partially solvent accessible, **OSF** repository: https://osf.io/52bng/). We noticed that the dataset of the *vicinal* cysteines also included cases in which the two cysteines are consecutive in the sequence (**OSF** repository: https://osf.io/52bng/). Disulfide bonds between sequence-adjacent cysteines are rare, but they can occur in proteins with either a *cis* or a *trans* peptide ^32^. Depending on the local structure, some of these pairs might undergo the proposed population shift mechanism without pronounced consequences, such as local misfolding. In contrast, for others, the *S-*nitrosylation could have major consequences on the local 3D structure. The dataset with sequence-adjacent vicinal cysteines could be interesting for future investigation with enhanced sampling simulations to account for an accurate description of changes in the population of folded/unfolded states upon *S-*nitrosylation.

An important parameter in the analysis of cysteine sites prone to redox modification is the residue pKa, which, in the case of cysteine residues in protein, has been estimated at around 8.5 in an unperturbed scenario ^33^. Reactive cysteines often have lower pKa values due to their protein environment and surrounding residues, which could contribute to stabilizing the thiolate. On the contrary, a hydrophobic or electronegative environment would raise the pKa by destabilizing the negatively charged forms. Nevertheless, this is an oversimplified view of cysteine reactivity at the cellular level.^33^ Marino et al., in a survey on PDB structures, found that cysteines that are targets for *S*-nitrosylation have higher pKa values (average 9.1) than the ones of redox cysteines.^34^ In general, pKa prediction alone is not a good proxy for discriminating SNO from non-SNO sites ^3,34^ In our dataset, we found that the SNO sites with a *vicinal* or *proximal* cysteine have a broad distribution of predicted pKas (**Figure 5C**). Some cases included SNO sites with very high pKa, suggesting an unsuitable environment for *S*-nitrosylation. It might be useful to explore pKa predictions further and perform them over an ensemble of conformations representative of the protein dynamics ^35^, as we did in the next section.

Finally, we evaluated whether the cases of proximal and vicinal cysteines were mutually exclusive. This turned out not to be the case, pointing out 88 proteins for which both cases of vicinal and proximal cysteines can be found around the known SNO sites (**Figure 5D**). Moreover, the dataset is equally divided between cases with a single SNO site characterized for the target protein and cases with multiple cysteines that undergo *S-*nitrosylation in the same protein (**Figure 5E**).

### Proximal cysteines in human S-nitrosylated proteins

*S-*nitrosylation is relevant in different diseases, including cancer ^36,37^. We thus investigated the 95 cases of SNO sites in 44 human proteins with proximal cysteines in more detail. SNOfinder analyzes static structures and uses relatively crude distance-based criteria to search for potential proximal cysteine candidates. On this subset of human proteins, we thus more carefully evaluated the reciprocal orientation of the two cysteines. Also, for the cases with proximal cysteines, we included the possibility of accounting for conformational changes, which could bring the residues closer in space using a coarse-grained molecular simulation approach (see Materials and Methods).

We collected coarse-grained simulations and reconstructed the full-atom models for the 44 proteins selected for simulations (**Figure 6/Table S3**). Enrichment analyses of these proteins against the Elsevier Pathway collection show a signature related to genes with mutations in cancer metabolic reprogramming (p-value < 0.005, odds ratio = 91.9). Moreover, enrichment analyses against ClinVar gene sets show signatures for hepatocellular carcinoma, lung cancer, glioma, and acute myeloid leukemia (p-value <0.05).

**Figure 6.**
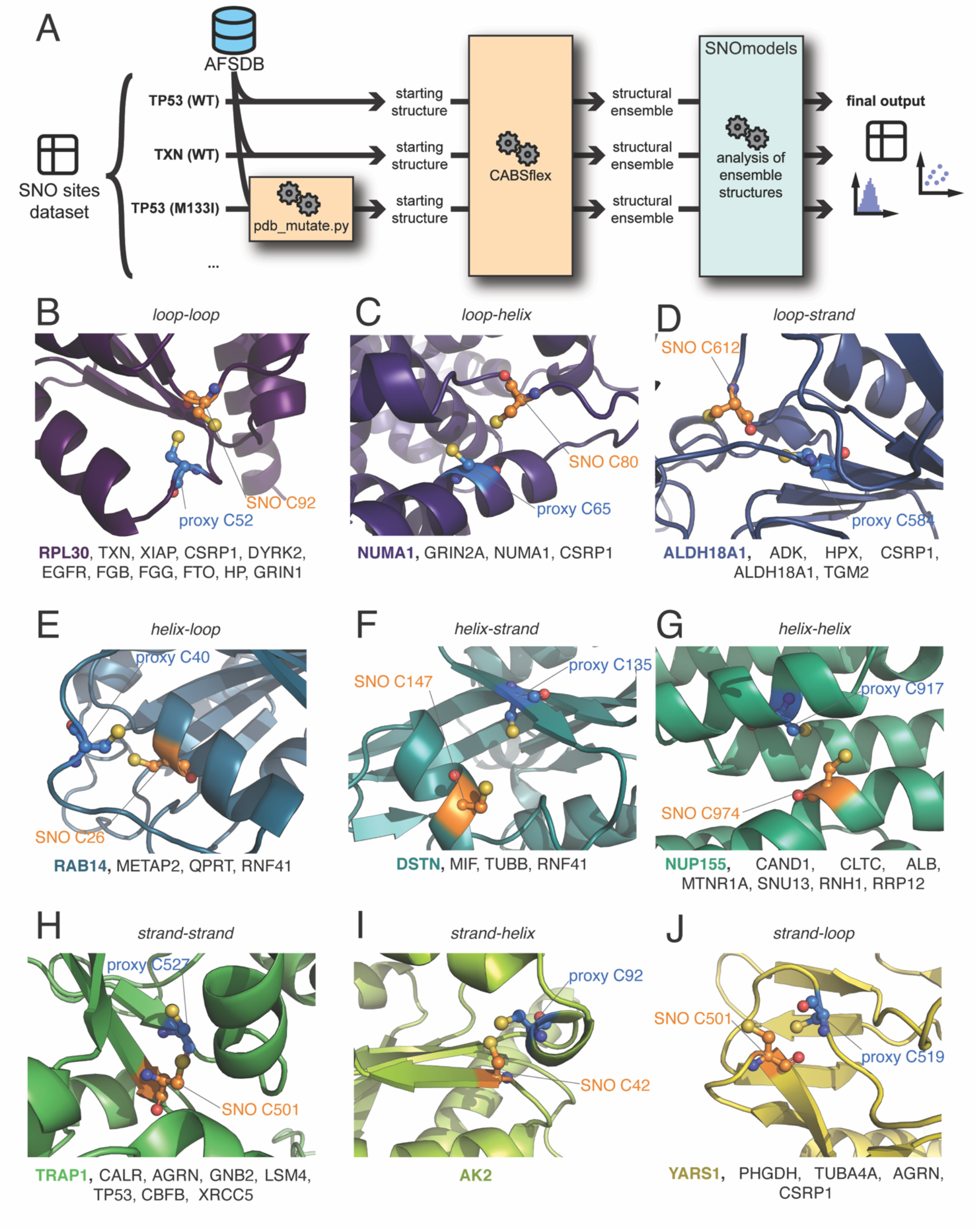
Analysis of reciprocal orientation and position of SNO and proximal cysteines in human *S*-nitrosylated proteins. A) Schematic representation of the workflow we used to generate and analyze structural ensembles for our SNO proteins having proximal cysteine residues. We start from the manually curated list of SNO sites with proximal cysteine sites for each protein, which was derived from the output of the SNOfinder pipeline (see Materials and Methods and Figure 5). For each protein, we obtained its model from the AlphaFold Model Structure Database and use CABS-flex to generate a structural ensemble for each. For p53, we also mutated one amino acid according to the mutation list we obtained for this protein (see Materials and Methods) and generated their own structural ensemble. Each structural ensemble included 20 models. We then used the SNOmodels pipeline to calculate relevant structural measurements for each model, such as dihedral angles and distances or interest, and we estimate the pKa of cysteine residues of interest as well as their relative access surface area (see Materials and Methods). Finally, we generated distribution and scatter plots, and calculated mean values and standard deviation for each measurement. B-J) We classified the identified proteins into nine categories depending on the secondary structural elements (strand, helix, loop) on which the SNO and the proximal cysteines are located: B) *loop-loop*, C) *loop-helix*, D) *loop-strand*, E) *helix-loop*, F) *helix-strand*, G) *helix-helix*, H) *strand-strand*, I) *strand-helix*, J) *strand-loop*. The cartoon representations show an example of a protein from each class (RPL30, NUMA1, ALDH18A1, RAB14, DSTN, NUP155, TRAP1, AK2, and YARS1, respectively). The spheres indicate the atoms of the *S*-nitrosylated (SNO, orange) site and the proximal cysteine (proxy, blue). At the bottom of each panel, we reported the list of the proteins classified in each class. We identified multiple pairs of cysteines for each class in different proteins apart from the *strand-helix* class, for which we identified only a pair of cysteines in AK2.

We then analyzed the simulations using another workflow, i.e., SNOmodels. The SNOmodels workflow calculates a relevant set of dihedral angles and distances corresponding to those used as collective variables in metadynamics (see above). SNOmodels also collects predicted pKa values and solvent accessibility for SNO-site and proximal cysteine for every model available in the 44 structural ensembles (**Figure S3, Figure 6A**).

As expected, we found TRAP1 cysteines in the middle domain as candidates from this screening. At first, we evaluated if the distribution plots obtained by the coarse-grain approach could qualitatively recapitulate what was observed for TRAP1 with the all-atom enhanced sampling presented in Figure 3 (**Figure S3)**. Interestingly, we observed that this simple approach could identify distance distributions and preferences for the side-chain dihedrals of the two cysteines close to what was observed with enhanced sampling for the reduced form of the TRAP1 middle domain.

The *S-*nitrosylated cysteine of adenosine kinase (ADK), methionine aminopeptidase 2 (METAP2), cullin-associated NEDD8-dissociated protein (CAND1), clathrin (CLTC), destrin (DSTN), E3 ubiquitin-protein ligase X-linked inhibitor of apoptosis (XIAP), FTO, QPRT, SNU13, NUP155, ALDH18A1, RAB14, RRP12, SPART, YARS1, XRCC5, have been identified in a high-throughput study with resin-assisted capture techniques coupled with iTRAQ labelling.^38^ In addition, the dual specificity tyrosine-phosphorylation-regulated kinase 2 (DYRK2), LSM4, MIF, CBFB, and RNF41 have been identified in a study with high-density protein microarray chips.^39^ Other candidates were also found from high-throughput experiments, as well as they have been reported in more in-depth investigations.

In the following, we divide the candidates into nine different categories depending on the secondary structural elements of the SNO and proximal sites (loop-loop, helix-helix, strand-strand, loop-helix, helix-loop, loop-strand, strand-loop, strand-helix, helix-strand).

We identified ten proteins classified as *loop-loop* (**Figure 6B**) including thioredoxin (TXN), XIAP, cysteine and glycine-rich protein 1(CSRP1), dual specificity tyrosine-phosphorylation-regulated kinase 2 (DYRK2), epidermal growth factor receptor (EGFR), fibrinogen beta chain (FGB), fibrinogen gamma chain (FGG), FTO, haptoglobin (HP), G protein-regulated inducer of neurite outgrowth (GRIN1), and 60S ribosomal protein L30 (RPL30).

As examples from this class, TXN is the prototype of disulfide-bridge formation between an SNO site and its vicinal cysteine ^14,15^. We also identified a pair of candidates for the formation of disulfide bridges of an SNO site (C73) with a proximal cysteine (C32) (**Figure 7A**). C32 is buried in the initial AlphaFold model. In contract, this residue can also explore more solvent-accessible conformations during the simulations due to the location of both cysteines in flexible loop regions, with Sγ-Sγ distances in the range of 3-7.5 Å (**Figure 7A**). We also noticed that the SNO sites and proximal cysteines seem to favor the minus state for their χ_1_ dihedrals, suggesting that the proximal cysteine can act as a sensor for the *S-*nitrosylation (**Figure 7A**). *S-*nitrosylation of C73 is also essential for interacting with procaspase-3, which can be trans-nitrosylated by thioredoxin ^40,41^. Our results suggest a new possible role for the redox modification of C73 in promoting intramolecular disulfide bonds with C32.

**Figure 7.**
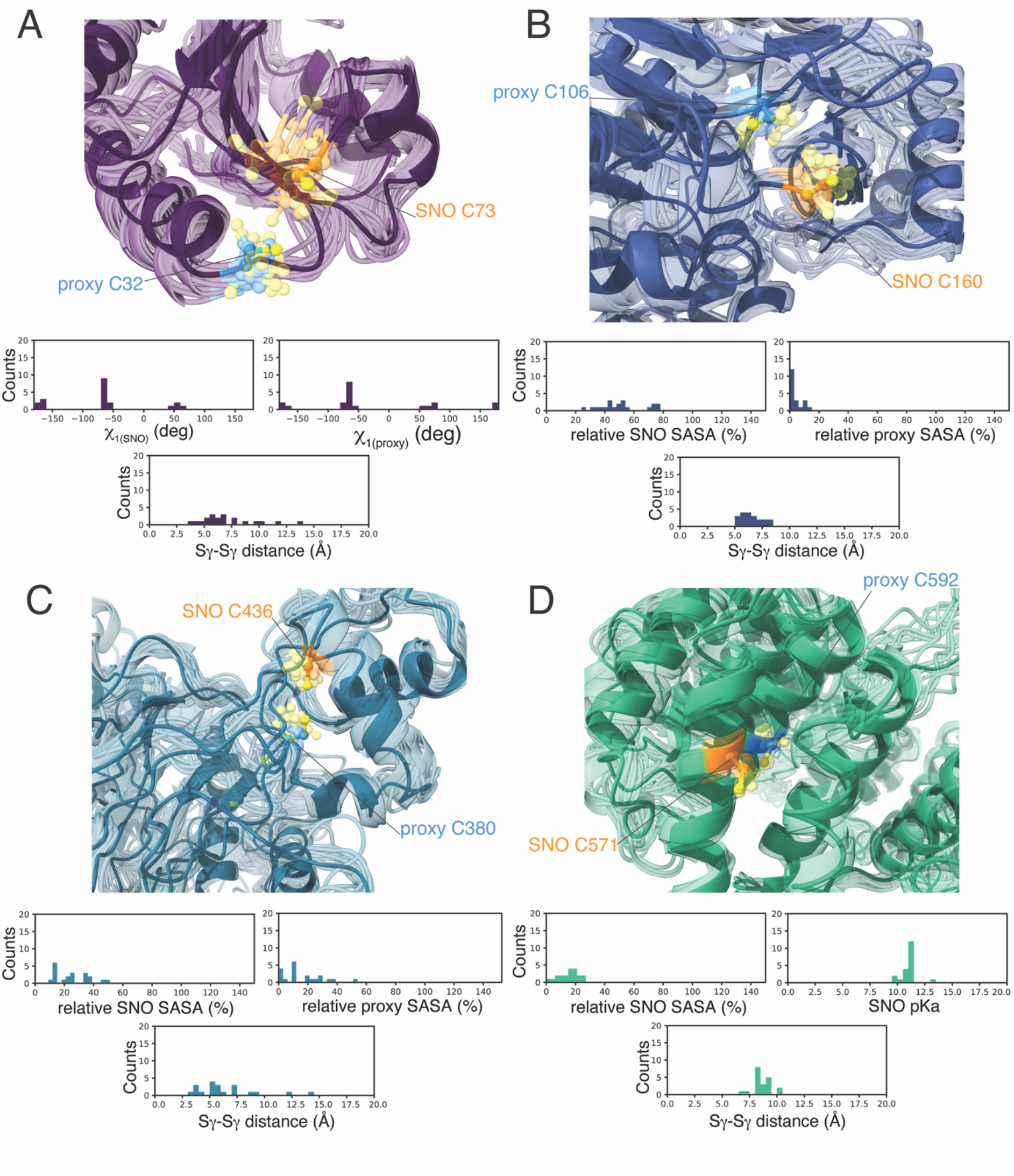
*S*-nitrosylated cysteines can induce conformational changes in the proximal cysteines in different human proteins. To account for conformational changes of the SNO and proximal cysteines, we collected coarse-grain molecular simulations of the 44 human proteins identified by the SNOfinder pipeline (see Materials and Methods). A-D) The cartoon show the structural ensembles calculated for A) thioredoxin (TXN, *loop-loop* class), B) adenosine kinase (ADK, *loop-strand* class), C) methionine aminopeptidase 2 (METAP2, *helix-loop* class), and D) cullin-associated NEDD8-dissociated protein (CAND1, *helix-helix* class). The spheres indicate the atoms of the *S-*nitrosylated (SNO, orange) site and the proximal cysteine (proxy, blue). The distribution plots at the bottom of each panel show the values calculated in the coarse-grain ensembles for i) χ_1(SNO)_, χ_1(proxy)_ and Sγ-Sγ distance (panel A), ii) relative solvent accessible surface area (SASA) of the SNO and proximal cysteines and Sγ-Sγ distance (panel B and C), iii) χ_1(SNO)_, pKa of the SNO cysteine and Sγ-Sγ distance (panel D). We calculated these values using the SNOmodels pipeline (see Methods). We observe that during the simulations, the SNO site C32 of TXN (panel A) and C160 of ADK (panel B) can undergo conformational changes and explore more solvent-accessible conformations, with Sγ-Sγ distances mainly in the range of 3-7.5 Å and 5-8 Å, respectively. Furthermore, the SNO sites and proximal cysteines of TXN seem to prefer the minus state for their χ1 dihedrals. The proximal C380 of METAP2 (panel C) is on a highly dynamic loop and assumes different orientations. On the other hand, the two cysteines of CAND1 (panel D) are in α-helices that constrain their positions, with Sγ-Sγ distances larger than 7.5 Å.

XIAP is a multi-functional E3 ubiquitin-ligase involved in the regulation of caspases but also inflammatory signals, mitogenic kinases, and cell proliferation, invasion, and metastasis ^42,43^. XIAP contains three baculoviral IAP repeats (BIR) motifs of approximately 70 residues. These motifs have been reported to be S-nitrosylated without major effects on XIAP ligase activity but compromising its anti-caspase-3 and antiapoptotic functions ^44^. In a previous work, cysteines 90, 213, and 327 of XIAP were found as S-nitrosylated ^45^. From our analyses of the data reported in dbPTM, we found C303 as a possible S-nitrosylated site with C327 as its proximal cysteine. Moreover, C327 and C300 are reported as SNO sites with SNOfinder with solvent accessibility lower than 10%. For this reason, they have been discarded from our downstream analyses. The two cysteines are in two loop regions and can reach distances as low as 3 Å in the simulations. However, the two cysteines, together with C300, coordinate one zinc ion, according to the analysis with AlphaFill ^46^. Cofactors are not included in the models deposited in the AlphaFold database. Therefore, it is likely that the *S-*nitrosylation of one of the two cysteine sites could cause major structural consequences for the zinc-binding site and cause misfolding. This is an interesting case study to address the interplay between *S-* nitrosylation and propensity for proteasomal degradation or other clearance mechanisms of misfolded proteins, similar to what was observed for the *S-*nitrosylated variant of TRAP1^7^. Moreover, *S-*nitrosylated XIAP has been detected in the brain of patients with neurodegenerative disorders ^47^, addressing the underlying structural mechanisms of its redox modification even more critical. The two cysteines (C373 and C347) are located on two flexible loops, and their sulfur atoms can get to distances as small as 2 Å (**Table S3**).

We found three cases classified as *loop-helix*, i.e., glutamate receptor GRIN2A, nuclear mitotic apparatus protein 1 (NUMA1), and CSRP1 (**Figure 6C**). NUMA1 is one example of a loop-helix class where the SNO site (C80) is on a long disordered loop, while the proximal C65 is placed on an α-helix.

The *loop-strand* class includes proteins with the SNO site located in a disordered region and the proximal site on a β-strand (**Figure 6D**). As examples of this class, we identified the adenosine kinase (ADK), hemopexin (HPX), CSRP1, ALDH18A1, and the protein-glutamine gamma-glytamyltransferase 2 (TGM2).

ADK has an SNO site (C160), which is partially exposed to the solvent in the initial model (SASA=21.7%). C160 is located in a loop region upstream of a short turn. This region undergoes substantial dynamics during the simulations and allows C160 to explore more solvent-accessible conformations (**Figure 7B**). In ADK, the proximal site (C106) is located in a β-strand, is more constrained regarding conformational changes, and is mainly buried **(Figure 7B**). The Sγ-Sγ distance ranges between 5-8 Å, with conformations resembling the ones observed for TRAP1 in the *S*-nitrosylated form. We observed a similar scenario for TGM2. These cases suggest that the *S-*nitrosylation could influence the side-chain conformation of the proximal cysteines, as well as affect the dynamics of the disordered regions where the SNO site is, introducing a variation to the mechanism observed for TRAP1. HPX includes an SNO site (C366) in a loop connecting two β-strands and a proximal cysteine (C408) on a β-strands belonging to a different β-sheet. In the AlphaFold model, the two sulfur atoms are predicted at a distance already compatible with disulfide bridge formation (around 2 Å). Still, according to the simulations, this pair can be found at distances from 5 to 10 Å in the reduced form.

The SNO site of ALDH18A1 (C612) is in a mobile loop that could undergo opening/closing motions interesting the proximal cysteine (C584) placed at the end of a β-strand of a parallel β-sheet. Nevertheless, we did not observe the two cysteines at distances lower than 7 Å in the coarse-grain simulations (OSF repository: https://osf.io/52bng/), suggesting that further exploration with enhanced sampling approaches is needed to understand this case in detail. The *helix-loop* class included methionine aminopeptidase 2 (METAP2), nicotinate-nucleotide pyrophosphorylase (QPRT), Ras-related protein Rab-14 (RAB14), and RNF41 **(Figure 6E**). METAP2 has the SNO site (C436) on an α-helix and the proximal site (C380) on a highly dynamic loop (**Figure 7C**). The high flexibility of C380 makes it a potential sensor of the modified variant of C436 upon *S*-nitrosylation. METAP2 is also an interesting drug target in cancer and other diseases.^48^ Moreover, METAP2 has been recently proposed as a target for PROTAC (Proteolysistargeting chimera) molecules^49^. In this context, the identification of METAP2 as interested by the population-shift mechanism induced by SNO could be the focus of further studies to evaluate if its *S*-nitrosylation is linked to increased proteasomal degradation, as previously observed for TRAP1 ^7^.

QPRT also belongs to the helix-loop class with the SNO site (C96) in the proximity of a bent N-terminal region of a long α-helix. The bending observed in the model is likely to be promoted by G100 and could contribute to conformations that bring C96 closer to the proximal cysteine C69, in agreement with the fact that in the simulations, we recorded distances lower than 4 Å. RAB14 has a proximal cysteine (C40) on a flexible loop which could undergo conformational changes that would bring its side chain closer to the SNO site (C26). The dynamics observed in the simulations suggest that the two residues can get closer than 5 Å in most of the simulation structures. Of note, in RAB14, the analysis with AlphaFill suggests that C26 interacts with GDP or GTP, which has not been accounted for in our simulations. Further studies with all-atom simulations will be needed to verify the influence of these interactions on the mechanism discussed here.

The *helix-strand* included pairs of SNO site and proximal cysteines from four different proteins, i.e., destrin (DSTN), macrophage migration inhibitory factor (MIF), tubulin beta chain (TUBB), and RNF41 **(Figure 6F**).

DSTN has a depolymerizing function on actin and is important for cell motility and cytoskeleton remodeling. The SNO site (C147) lies on an α-helix, and its proximal cysteine lies on a β-strand in front of it. This arrangement likely limits how close these two residues can get, meaning that the two side chains can reach distances around 5 Å only in a small number of ensemble structures. Similarly, the macrophage migration inhibitory factor (MIF) has a structural arrangement for the pair of cysteines in which the SNO site is located on an α-helix placed on the top of a β-sheet which includes the proximal cysteine in a central position. In this case, structural constraints might also be at play, with conformations at a distance rarely lower than 7 Å.

The SNO site of TUBB (C239) is at the C-terminal extremity of an α-helix in a bent region. This suggests a certain degree of flexibility which can contribute to conformations where the site can reach the proximal cysteine, confirmed by the simulations.

Overall, the helix-strand class poses structural constraints which could make some of the targets in this category unlikely candidates for the disulfide-bridge formation and the population-shift mechanism. Alternatively, the proteins can still undergo the SNO-induced mechanisms with major consequences on their structural integrity.

Eight protein candidates have been found associated with the *helix-helix* class, i.e., CAND1, CLTC, ALB, MTNR1A, SNU13, NUP155, RNH1, and RRP12 (**Figure 6G**). In addition, ALB included multiple SNO sites with this arrangement.

CAND1 is an F-box protein exchange factor, which is coupled with cycles of neddylation. In the deneddylated state, CAND1 can bind CUL1-RBX1 and promotes the dissociation of the SCF (SKP1-CUL1-Fbox) complex and the exchange of the F-box protein.^50^ CAND1 is important in the recycling process of the cullin scaffold. It has been proposed that cells lacking CAND1 cannot easily adapt to handle large pools of F-box proteins.^51^ In this case, the two cysteines are in α-helices facing each other. In the simulations, we did observe cys-cys distances larger than 7.5 Å. This suggests that the folded helices pose constraints on the contacts between the two cysteines, emphasizing the importance of including dynamics in the study of the candidates from the SNOfinder screening.

CLTC is known for its role in clathrin-mediated endocytosis but also participates in several other cellular functions. CLTC has also been described as a molecular shape-shifter for its ability to assemble into different polyhedral lattices or tubular structures ^52^. The two cysteines are located in two facing α-helices. The reciprocal orientation of the helices seems to accommodate the possibility for opening/closing motions, with the consequence of allowing conformations where the two cysteines get at distances of 4 Å from each other (**Figure 7D**). CLTC is also one of those cases in which its conformational dynamics can influence the SNO site pKa and promote conformations where the cysteine is more likely to be reactive (**Figure 7D**).

NUP155 is a protein of the nuclear pore and has a helix-helix interaction for the SNO site C974 and its proximal cysteine C917, which in the simulations can reach distances as low as 3 Å. RNH1 is a ribonuclease inhibitor in which the *S*-nitrosylable cysteine (C75) and its proximal partner (C45) are in the middle of two α-helices facing each other, and conformational changes of their side chains could promote orientations prone to disulfide bridge formation, featuring distances in the range of 3.5-8 Å according to the simulations. The proximal cysteine remains buried during the simulations where the SNO site can explore mostly partially or fully solvent-exposed conformations.

RRP12 has a SNO site (C317) at the end of an α-helix, and its location suggests a region that could also explore more disordered conformations connecting the α-helix to another adjacent α-helix. This suggests that RRP12 could also be classified as a loop-helix target, depending on the conformational heterogeneity observed for this protein. On the contrary, the proximal C261 is in the middle of a well-folded α-helix. In this case, we observed distances often higher than 7 Å during the dynamics, pointing out RRP12 as a candidate for further studies with enhanced sampling to understand the structural constraints in the area where the cysteines are located. A similar scenario also applies to MTNR1A.

Summing up, as observed for the helix-strand class, when two cysteines are both located in α-helices, the 3D architecture of the target protein can pose structural constraints that do not allow for conformations at a distance compatible with the mechanism proposed in this study or which is at play would have major consequences on the native protein structure.

In the *strand-strand* class, together with TRAP1, we found CALR, AGRN, GNB2, LSM4, TP53, CBFB, and XRCC5 (**Figure 6H**).

GNB2 is also classified as a case of strand-strand pair of cysteines. However, this case does not constitute a suitable candidate for the population shift mechanism proposed for the interaction between the SNO site and its proximal cysteine due to structural constraints posed by a β-strand located between the two cysteine sites.

The pair of cysteines in LSM4 is located in two different two-stranded small β-sheets bent towards each other and connected by short disordered regions. The disordered regions could serve as a hinge for opening and closing movements that could bring the two cysteines at a closer distance.

Another example of this class is CBFB, discussed in detail in the next section, as a cancer driver.

In XRCC5, the two cysteines are located on different elements of a small barrel structure. The SNO site (C339) is at the end of a short β-strand. Its location suggests that conformational changes of its backbone or sidechain dihedrals could promote conformations that could face the proximal cysteine (C249), which is in the middle of a β-strand. This is confirmed by the simulations where the two residues are found at a distance below 4 Å in almost half of the structural ensemble.

Adenylate kinase 2 (AK2) is the only case among the 44 protein candidates belonging to the *strand-helix* class. In the Alphafold model, the SNO site of AK2 (C42) is located in a β-sheet in the protein core and faces a proximal cysteine (C92) on a small 3.10 helix at a distance of approximately 2 Å (**Figure 6I**). In the simulations with the reduced cysteines, the residues remain at short distances in most of the ensemble structures, and the SNO site fluctuates between partially and more solvent-exposed states lowering its pKa and improving its reactivity.

The *strand-loop* group included PHGDH, YARS1, TUBA4A, AGRN, and CSRP1 **(Figure 6J**). D-3-phosphoglycerate dehydrogenase (PHGDH) and YARS1 depending on the structural heterogeneity of the cysteine sites, can be seen as strand-loop or loop-loop classes. This is due to the collocation of their SNO sites at the very end of a short β-strand.

TUBA4A is another example of a strand-loop target where there might be structural constraints that do not allow the SNO site and the proximal cysteine to get closer than the distance observed in the AlphaFold model (i.e., 7.8 Å). In this case, the simulations suggest that the sites can be more flexible than what was observed in the initial structure, exploring conformations with an Sγ-Sγ distance in the range of 3-7 Å which fits well with the structural mechanism proposed in this study, enforcing the notion of the importance to account for protein dynamics.

We also identified three proteins that include multiple reported S-nitrosylations and multiple candidates as proximal cysteines for each SNO site (AGRN, ALB, CSRP1). AGRN has a SNO site (C405) with two possible candidates as proximal cysteines (C395 and C414), belonging to the strand-loop and strand-strand classes, respectively. EGFR also has two possible proximal cysteines (C199 and C207) for the *S*-nitrosylable residue C190. In both cases, they are classified as belonging to the loop-loop category. ALB featured 14 pairs of possible SNO sites and proximal cysteines, all belonging to the helix-helix class. CSRP1 has four different SNO sites, of which three have two possible proximal cysteines as candidates. The CSRP1 cysteine pairs cover various classes, mostly belonging to the strand-loop category and with cases of loop-strand, loop-helix, and loop-loop classes. Haptoglobin (HP) has two possible SNO sites with one proximal cysteine, each belonging to the loop-loop class, both at distances compatible with disulfide bridge formation. SNOfinder identifies two cysteines that can be both *S*-nitrosylated (C798 and C744) and serve as proximal cysteines to each other in the Glutamate receptor ionotropic, NMDA 1 (GRIN1), both belonging to the loop-loop class. TP53 has one SNO site satisfying the criteria of the SNOfinder protocol (C124), with two possible proximal cysteines belonging to the strand-strand class. It is described in more detail in the next section as a target of interest in cancer. RNF41 has one SNO site (C134) with two possible candidates as proximal cysteines belonging to the helix-strand (C116) and helix-loop class (C120), respectively (**Table S4**).

GRIN2A, CBFB, CALR, EGRF, and TP53 are discussed in more detail in the next section concerning their role as cancer driver proteins.

### Modification of population shift mechanisms induced by cancer mutations

Among the candidates with proximal cysteines that could act as sensors of the *S*-nitrosylation to promote disulfide-bridge formation relying on vicinal or proximal cysteine, the identified human proteins are of interest in relation to cancer. Indeed, GSNOR, the enzyme devoted to removing *S*-nitrosylation from proteins, is known to be downregulated in several cancer types^6,8,53^, suggesting that *S*-nitrosylation and its structural consequences could be enhanced in a cancer setting. We thus evaluated if any of these proteins are classified as an oncogene or tumor suppressor according to the COSMIC Cancer Gene Census ^54^.

We identified three tumor suppressors: CTNNA1 (C772 and vicinal cysteine C767), GRIN2A (C320 and proximal C87), and CBFB (C25 and proximal C124). We also found two oncogenes with a pair of SNO sites and proximal cysteines (C181 and vicinal C184 in HRAS, C105 and proximal C137 in CALR), as well as the oncogenic protein EGFR with multiple candidate cysteines (vicinal C194, and proximal cysteines C199 or C207) in the proximity of its SNO site (C190). We also found the guardian of the human genome, TP53, which is classified by COSMIC as a dual-role gene thanks to its properties as a tumor suppressor and oncogene. TP53 has a reported *S-*nitrosylation at C124 and two candidate proximal cysteines C135 and C141.

For each cancer driver protein with at least one proximal cysteine to the SNO site, we retained the residues in the surrounding of each SNO and proximal/vicinal sites as detailed in the Methods. In parallel, we collected known cancer variants or variants reported in ClinVar (see **Materials and Methods** for more details). For each dataset of cancer mutations in the candidate driver proteins, we retained the ones located in the surrounding of the cysteines as defined above for further analysis. At first, we evaluated if any of the mutations is expected to alter the structural stability of the protein using a structure-based framework recently proposed by our group^55^ (**Methods, Tables S5**). All those mutations that seem to have a neutral or uncertain effect on the structural stability of the candidate protein are interesting candidates for further analyses in connection with the population shift mechanism proposed here. We reasoned that the destabilizing mutations would mostly result in proteasomal degradation or other clearance mechanisms or aggregation at the cellular level, which would overshadow changes in the structural mechanisms promoted by *S-*nitrosylation.

In particular, we considered the possible alterations that the mutations could cause in connection with different effects: i) altering the propensity of the SNO site to be S-nitrosylated (either acting directly on the *S-*nitrosylable cysteine or changing the properties of the environment with consequences on the predicted pKa for the residue); ii) interfering with the population shift mechanism, for example reducing the accessibility to the *plus* or *minus* states of the χ_1_ of the proximal cysteine or the SNO site or altering the distance between the two sites.

In TP53, the cysteines are located in the DNA-binding domain (DBD), which is heavily mutated in cancer. Virtually, each residue of the DBD of TP53 can be targeted by mutations in cancer samples ^56^. We here used a dataset of mutations curated in our previous work ^56^ and further, included variants from ClinVar ^57,58^. This list also contains mutations on the SNO site and proximal cysteine (C124 to F, G, L, R, S Y, C135 to G, S, T, and C141 to A, S, Y). The residues in the surroundings of the SNO and proximal cysteine sites are S116, V122, T123, T125, M133, Q136, T140, P142, V143, Y234, N235, Y236, V272, V274, and P278. We identified 54 variants (see **Table S6**) at these positions predicted as neutral or uncertain in terms of changes in folding free energy that we could explore further in connection to the SNO-induced population shift mechanism. Thus, we carried out simulations with the same coarse-grained approach applied for the reduced variant of TP53 for each of the 54 mutations and analyzed them with the SNOmodels pipeline.

The SNO-site (C124) is at the bottom of a DNA-binding loop, L1, where a key residue (K120) for interaction with the DNA is located. This loop is known to undergo conformational changes from recessed (optimal for DNA binding) to more extended states. It is likely that the modification of the cysteine could influence the conformational changes of the loop, with consequences on DNA binding. The two candidate proximal cysteines reside at the extremity of a small loop (135-141), and the three cysteines form a tri-partite cluster of SH groups in the 3D structure, suggesting the possibility of a mixture of redox states upon post-translational modification (**Figure 8A**).

**Figure 8.**
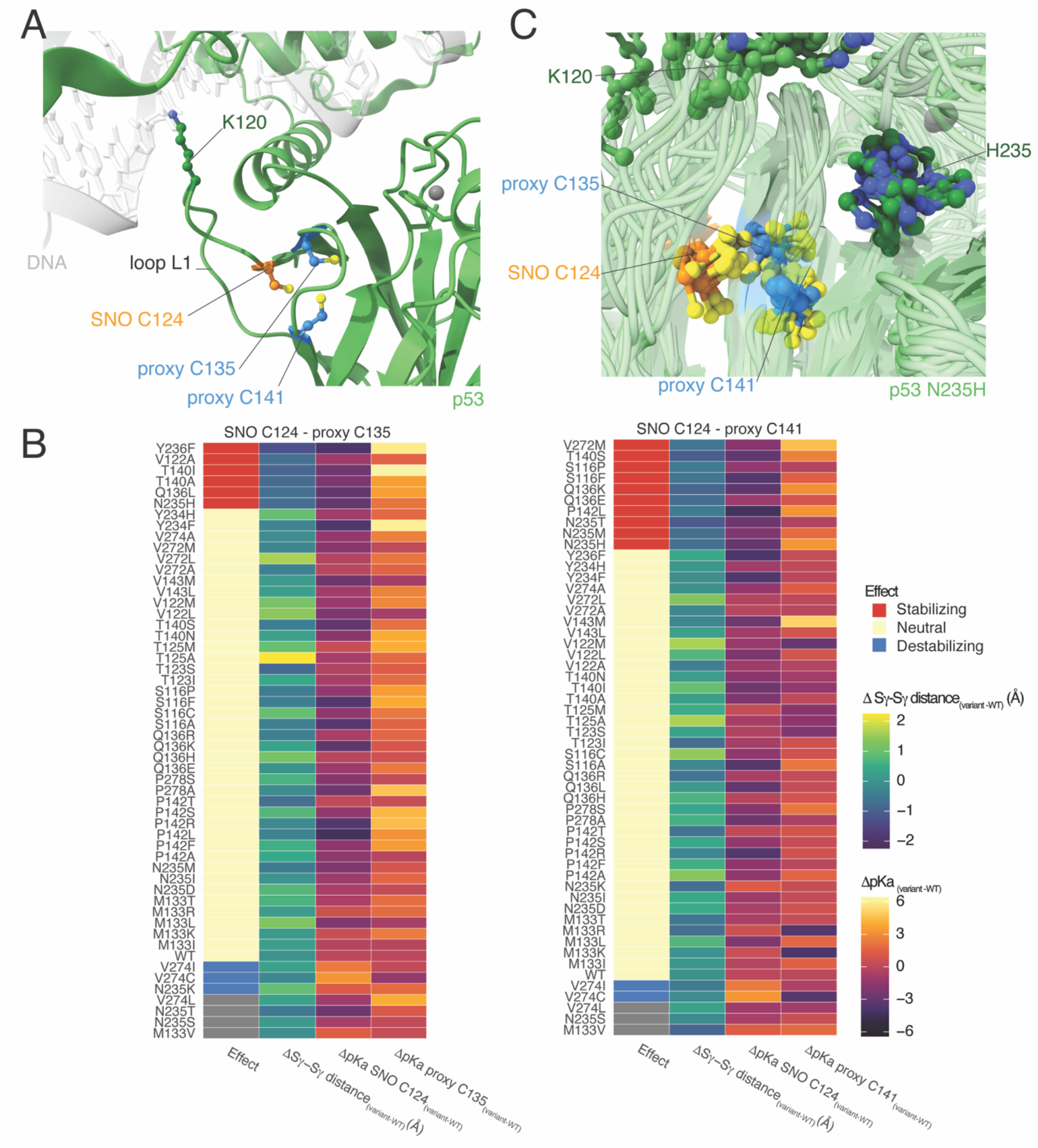
Cancer variants in TP53 could modify the population-shift mechanism induced by *S*-nitrosylation. A) The green cartoon shows the AlphaFold model of the tumor protein 53 (p53). We show, as a reference, the location of the DNA from the X-ray structure of the p53 tetramer (PDB entry 3KZ8) as a grey transparent cartoon. We used Alphafill to include the missing zinc atom (grey sphere) in the DNA-binding domain of p53. The stick and ball representation highlight the *S*-nitrosylated site C124 (SNO, orange), proximal C135 and C141 (proxy, blue), and K120 (dark green), a key residue for interaction with DNA located in the DNA-binding loop L1. B) We analyzed the coarse-grain simulations of the variants of p53 in the proximity of the SNO site and proximal cysteines and compared them to the wild-type (WT) p53. The heatmaps summarized the values calculated for the SNO site C124 and proximal C135 and C141 (left and right heatmap, respectively). Each heatmap shows the distance between the two sulfur groups of the two cysteines (Sγ-Sγ distance, second column) and the predicted pKa of the SNO site and proximal cysteines (third and fourth column, respectively). We then classified each variant (first column of the heatmaps) on its effects on the structural mechanisms of *S*-nitrosylation. We classified the variants as i) stabilizing (decreased pKa of the SNO site and shorter Sγ-Sγ distance respect to WT), ii) neutral (similar pKa of the SNO site and similar Sγ-Sγ distance between variant and WT) and iii) destabilizing (increased pKa of the SNO site and longer Sγ-Sγ distance respect to WT, and cases with a large increase of pKa of the SNO site). Stabilizing variants are highlighted in red, while blue indicates the destabilizing ones. Grey tiles indicate unknown/uncertain classification. We observed several variants of p53 that are classified as stabilizing for the SNO site C124 and proximal C135 (V122A, Q136L, T140I, T140A, N235H, Y236F) and C141 (S116F, S116P, Q136E, Q136K, T140S, P142L, N235H, N235M, N235T, V272M). These variants could make the protein environment more favorable to the population-shift mechanism or *S*-nitrosylation. On the other hand, we observed few variants that could have destabilizing effects and impair the structural mechanisms (V274I, V274C, N235K for proximal C135 and V274I, V274C for proximal C141). C) The cartoon representations show the structural ensemble of the coarse-grain molecular simulations of the stabilizing variant N235H. The stick and ball representations highlight the *S*-nitrosylation site C124 (SNO, orange), proximal C135 and C141 (proxy, blue), K120 and H235 (dark green). We observed that N235H favors shorter distances between the SNO site C124 and the proximal C141, Cβ-Sγ-Sγ-Cβ dihedral values around −90 degrees, and lower pKa of C124. These effects suggest that N253H could enhance the *S-*nitrosylation of C124 and the population-shift mechanism for disulfide-bridge formation.

We analyzed the coarse-grain simulations of the mutant variants in the proximity of the SNO site and proximal cysteines in terms of different parameters (i.e., Sγ-Sγ distances, Cβ-Sγ-Sγ-Cβ dihedral angle, χ_1_ side-chain angles, predicted pKa, and solvent accessibility, **Table S6**). We then classify these variants according to the following criteria for each pair of SNO and proximal site: i) no effects on the structural mechanisms related to *S*-nitrosylation in the case the results were similar between wild-type and mutant variant; ii) destabilizing mutations that impair the mechanism in the case the mutations induced an increase of pKa of the SNO site and increased distance between the cysteines, as well as cases with large increase of pKa; iii) stabilizing mutations that could make the environment even more favorable to the population-shift mechanism or *S*-nitrosylation (decreased pKa and shorter Sγ-Sγ distance). The results are summarized in **Figure 8B** and detailed in **Table S6**. Most variants with destabilizing effects act on the pKa directly, increasing its value and, thus, making it less prone to *S*-nitrosylation.

As an example of stabilizing mutation, N235H seems to favor shorter distances between C124 and C141, the exploration of structures with Cβ-Sγ-Sγ-Cβ dihedral values of −90 degrees, as well as to lower the pKa of the SNO site. This effect seems to be triggered by the histidine side chain that tends to point toward C141 and its sequence-adjacent residues in the mutant variant. In the wild type variant, the asparagine points in the opposite direction, away from the proximal cysteine. Overall, these features suggest N235H is an enhancer of the *S*-nitrosylation of C124 and the population-shift mechanism to induce disulfide-bridge formation (**Figure 8C**). In addition, T140A is also predicted with a stabilizing effect on the SNO-induced mechanism (**Table S6**) and annotated in ClinVar as a variant of uncertain significance (**Table S5**). Considering that, in cancer cells, *S*-nitrosylation is deregulated and prominent, the stabilizing effect on TP53 redox modification promoted by T140A constitutes a potential driver mutation for future studies.

## Discussion

Cysteine, via the sulfhydryl group, plays a myriad of different functional roles, including the capability to bind metals, electron donation, redox catalysis, and redox modifications^59^. The frequency of cysteines in the proteomes of modern species has been increasing^60^ attesting its functional importance. Cysteines seem to be conserved especially when they are associated with clusters with other cysteines in the 3D structure^61^. Cysteines are also the only residues that can form covalent bonds through their side chains, known as disulfide bonds or bridges. Disulfide bridges can also alternate between reduced and oxidized forms in mature proteins ^62^ and can be promoted by redox modifications, such as *S-*nitrosylation ^63,64^. Disulfide bridges should thus be seen as not “locked in” in biological systems ^33^.

In this work, we unveiled a new mechanism for structural changes induced by *S-*nitrosylation. Apart from already known cases of vicinal cysteines where *S-*nitrosylation of one of the two sites can promote disulfide bridge formation, we here identified cases in which the nearby cysteine is not close in sequence, but it is the 3D structure. We have shown that the intrinsic dynamics of these proteins can explore conformations prone to disulfide bridge formation and that such states seem to be favored by the *S-*nitrosylation of one of the two cysteines. We refer to this mechanism as population shift induced by *S-*nitrosylation. We reported a full investigation for TRAP1, the first case where we identified the mechanism, including experimental support to our findings. We then expanded our knowledge of the population shift mechanism with a high-throughput study, starting with the curation of known *S-*nitrosylated cysteines. This suggested that the identified mechanism might be more general, including a range of cases in which the specifics on how the *S*-nitrosylated and proximal cysteines come into contact vary, allowing us to define nine different structural classes. In the study, we focused on structural characterization of the protein candidates from the human proteome in connection to cancer, but the workflow here applied also provided a rich dataset to explore similar patterns in other species, along with the possibility to investigate more cases of vicinal and sequence-adjacent vicinal cysteines.

Ad added value of our work is also the development of a workflow based on two tools, i.e., SNOfinder and SNOmodels, which can be reapplied, in the future, to update the source of candidates with proximal or vicinal cysteines in parallel with new annotations available in dbPTM. Moreover, we noticed that the current annotations in dbPTM might not cover all the available *S-*nitrosylation reports in the literature. For example, we noticed that CAND1 *S-* nitrosylation was also provided by other proteomics studies ^65^. Our workflows can also be used on a list of manually curated *S*-nitrosylated sites, as far as the input format is coherent with the format of the dbPTM CSV files.

Moreover, the possibility to use a computationally inexpensive coarse-grained approach to model protein dynamics demonstrated the importance of structural investigations of the candidates, as it allowed us to rule out cases in which the structural constraints posed by the three-dimensional architecture of the protein might impair the population shift mechanism. For example, we observed that the protein architecture of some of the candidates in the helix-strand, helix-helix, and strand-strand classes restrained the movement of the SNO site or the proximal cysteine so that the formation of a disulfide bridge would be unlikely without major consequences on the structure. Examples of this class are DSTN, and MIF in the helix-strand class, as well as CAND1, RRP12, and MTNR1A in the helix-helix class. In addition, cases in which the cysteines are placed in β-strands could feature similar structural constraints (TUBA3A and GNB2).

In the future, the candidates provided here could become interesting targets for investigations with the enhanced sampling protocol proposed for TRAP1 to validate further the cases in which the population shift mechanism is at play. Moreover, the results from this work provide a rich ground to design simulations based on quantum mechanics-molecular mechanics or - molecular dynamics (QM/MM or QM/MD)^66^ or polarizable force fields^67^ to understand better the reactivity of *S*-nitrosylated cysteines with their protein environment, the mechanism of disulfide bridge formation and the effects associated with the high polarizability of the SNO group.

All-atom simulations and experimental assays will also be an interesting toolkit to apply to the hits identified in this study to understand if the proposed population shift mechanism induced by *S*-nitrosylation and consequent disulfide-bridge formation hold up for all the cases we proposed, and result in promoting misfolding, protein aggregation or other functional consequences. We know that, in the case of TRAP1, *S*-nitrosylation makes the protein more sensitive to undergo proteasomal degradation ^7^. However, we still do not know if this is: i) a direct consequence of *S*-nitrosylation or ii) induced upon disulfide bridge formation, or,actually, iii) inhibited when S-nitrosylation resolves into an SS adduct. How big is the proportion of *S*-nitrosylated *versus* SS derivatives in TRAP1, and how this ratio is affected by the cellular environment, or by the presence of enzymes which can interfere with the thiol redox state in proteins (e.g., glutaredoxins, nitrosylases or denitrosylases) are questions still unsolved. Based on results shown here and in our previous papers ^7,17^, we can just state that both these cysteine modifications are detected in TRAP1 and *S*-nitrosylation at C501 is induced both in vitro (i.e., exposure of recombinant TRAP1 to elevated fluxes of NO) and in vivo (i.e., in GSNOR-KO cells). Further investigations are needed to understand if disulfide-bridge may also act as a biologically functional modification of TRAP1, which takes place depending on the concentration of NO (or NO-releasing molecule, e.g., GSNO), as already proposed for OxyR^68^, a prokaryotic transcription factor acting as a prototype of this class of redox-sensitive proteins.

Due to the involvement of *S*-nitrosylation in neurodegenerative diseases^69^, the knowledge of the relationship between the proposed mechanism and propensity to protein misfolding goes beyond the mere interest of fundamental research and could have a great translational impact. Mutations in cysteine residues have been estimated to lead to genetic diseases more often than expected based on cysteine abundance ^70^. This makes the mechanism here proposed also more relevant from the point of view of biomedical research because disease-related mutations can alter the proposed mechanism, not only acting directly on the SNO site and its proximal cysteines but on the surrounding residues and changing the propensity for *S*-nitrosylation or the conformational preferences of the two sites that are important for the optimal formation of the disulfide bonds. In this context, we investigated mutations found in cancer studies for TP53 in the proximity of the *S-*nitrosylation site C124 and its proximal cysteines. In previous work, strategies to rescue TP53 by restoration of L1 loop conformation through cysteine modulations have been proposed, including C124 ^71–73^, including the development of drugs as PRIMA-1, which can rescue R175H TP53 mutants by targeting C124 (and C277)^74^, emphasizing the importance of unveiling structural mechanisms related to redox modifications of C124.

Our work identified three variants (N235K, V274C, V274I) that could destabilize the *S*-nitrosylation of C124 and/or the populations-shift mechanisms, mostly acting on the pKa of C124. Intriguingly, we also proposed ten variants with stabilizing effects resulting in a lowering of the pKa and shorten distances or favorable geometries for the population-shift mechanism, including mutations in the N235 site.

## Materials and Methods

### Analyses of TRAP1 experimental structures deposited in PDB

We used PDBSWS^75^ to retrieve structures, including coordinates for C501 (human TRAP1) and C516 (*Danio rerio* TRAP1) using the Uniprot ID Q12931 and A8WFV1, respectively. We then measured: i) the distance between the two sulfur atoms in the two cysteines, ii) the dihedral angle Cβ-Sγ-Sγ-Cβ, iii) the χ_1_ dihedrals for the SNO site (C501 and C516 in human and *Danio rerio* TRAP1, respectively) and the proximal cysteine (C521 and C542 in human and *Danio rerio* TRAP1, respectively).

We evaluated the geometries of the two cysteines using DSDBASE 2.0^76^, which classifies possible disulfide bridges from 3D structures in the PDB according to their stereochemical quality (A, B, C or D). ‘A’ refers to disulfide bridges with dihedral angles and S-S bond distances within the accepted range. ‘B’ is used for geometrically suitable pairs to form the S-S covalent bond but with distorted geometry. If the sites are too closed for a disulfide bridge, they are graded as ‘C.’ In addition, grade ‘D’ refers to pairs of residues that satisfy the distance criteria but with other issues in terms of geometric properties.

DSBBASE annotates all possible pairs of residues to accommodate a disulfide bridge without sterical constraints. If a pair of residues is not found, it does not satisfy the applied stereochemical parameters.

### Molecular dynamics simulations

The structures of TRAP1 from *Danio rerio* used in our previous work ^17^ in its reduced and *S*-nitrosylated form were used as starting structure for the simulation, retaining only the coordinates of the region corresponding to the middle domain (TRAP1_311-567_). We used this construct instead of the full-length protein since, in this work, we are interested in describing local effects induced by *S*-nitrosylation. Working on a smaller domain provided an advantage from the point of view of the computational time required to collect the enhanced sampling simulations. Metadynamics simulations, using the well-tempered ensemble approach^77^ were performed with GROMACS 4.6 ^78^ with the PLUMED 1.3 plugin ^79^ for 300 ns. The proteins were solvated in a dodecahedral box, and we applied periodic boundary conditions. The protocol and parameters to prepare the simulations followed the same approach applied to a previous case study ^80^. We used eight force fields (**Table S2**) to simulate the reduced variant of TRAP1_311-567_. We then selected ff99SB*-ILDN to compare the reduced and the *S*-nitrosylated forms. Moreover, we collected a one-μs unbiased molecular dynamics simulation of the oxidized form of TRAP1, where the two cysteines form a disulfide bridge, to be used as a reference of the values of the collective variables used in metadynamics. The simulations were carried out at 298 K in the canonical ensemble. We extended the simulations with the ff99SBstar-ILDN force field up to 500 ns to verify that the resulting free energy profiles did not change the overall conclusions of the work. The collective variables used are described in the results. More information on the parameters for the metadynamics simulations is reported in an OSF repository associated with the publication (https://osf.io/sfrkc/).

### SNOFinder and bioinformatic analyses of *S*-nitrosylated proteins from dbPTM

We have performed a comprehensive structural analysis to identify potential cysteine residues close to known *S*-nitrosylation sites. We have first downloaded a list of all available *S*-nitrosylation sites from dbPTM^4^. This dataset consists of 4172 SNO sites, annotated as belonging to a total of 2279 proteins from a variety of organisms (i.e., the 4172 sites were annotated with 2279 unique UniProt accession numbers or UniProt AC). We then cross-reference its contents with the AlphaFold Protein Structure Database (AFPSDB) by UniProt AC to identify all the proteins of the dataset that had an available AlphaFold structural model. Of the dbPTM dataset, 46 UniProt ACs were not found in the AFPSDB, corresponding to 100 *S*-nitrosylation sites. For each protein with an associated AlphaFold model, we then i) downloaded the corresponding model from the online repository, ii) used the PyInteraph2 software ^81,82^ to calculate a center of mass protein structure network with distance cut-off 8 Å, as an inter-residue contact map, iii) processed the resulting contact map to identify only cysteine residues that were in contact with known *S*-nitrosylation sites. We then annotated each S-nitrosylation site and its associated proximal cysteine residues with the associated AlphaFold pLDDT AlphaFold2 model quality score, relative solvent accessible surface area calculated with the NACCESS program ^83^ on the AlphaFold model, secondary structure assignment calculated using DSSP ^84^ and pK_a_ values predicted by PROPKA ^85^. Finally, for each entry, we calculated the distance between the sulfur atoms of the SNO site and the proximal cysteine, using MDAnalysis ^86^ to parse the AlphaFold model PDB files.

We started investigating 2279 proteins with 4172 known sites overall. We then removed from the analysis 46 proteins and 100 sites since we did not find a corresponding model structure in the AlphaFold database ^30^ and analyzed overall 2233 protein structures and 4072 SNO sites. We also found that 22 *S-*nitrosylation sites did not correspond to cysteine residues in the corresponding AlphaFold model and one case whose residue number was not included in the AlphaFold model. In addition, as a control, we repeated our analysis by increasing the PyInteraph2 distance cut offs to 11 Å.

The Snakemake workflow SNOFinder is available as a GitHub repository (https://github.com/ELELAB/SNO_investigation_pipelines), and the data produced for this study are deposited in an OSF repository https://osf.io/52bng/.

The enrichment analysis was carried out using EnrichR ^87,88^.

We discarded from the analyses RPL37A (UniProt entry C9J4Z3) since the reference provided by dbPTM did not refer to an article discussing *S-*nitrosylation. We also revised in **Table S3** the UniProt ID for METAP2, which is MAP2_HUMAN and not AMPM2_HUMAN as provided by dbPTM.

We also discarded from further analyses proteins that had too unstructured AlphaFold models or where the cysteine was located in regions with very low pLDDT score and long disordered loops or linkers (more than 20 contiguous disordered residues). For similar cases, more accurate and computationally intensive simulations would be required to properly assess the conformational changes of the SNO site and its proximal cysteines.

### CASB-flex simulations

For the 44 selected human targets, we collected simulations with CABS-flex ^89^. We used models from the AlphaFold database retrieved by SNOFinder and trimmed some of them to remove regions with low pLDDT scores, except loops shorter than 20 residues that connected folded domains. On each trimmed structure, we then ran a CABS-flex simulation performing 20 temperature annealing cycles, each consisting of 100 Monte Carlo cycles of 100 steps each, without setting any restraint. The only exception was the TP53 DNA binding domain (DBD). We used as starting structure a pre-processed AlphaFold model as described elsewhere [ref]. In the p53 CABS-flex runs, we decided to restrain the residues that make up the protein’s zinc ion (Zn^2+^) coordination site in the experimental structures binding site to ensure that its conformation would stay similar to one of the experimental structures, as CABS-flex doesn’t support cofactors such as zinc, to the best of our knowledge. In particular, we added distance restraints between each pair of the Cα atoms of residues 176, 179, 238, and 242, using as equilibrium distance for the restraints the distances present in an experimental, zinc-bound structure of p53 (PDB ID 2XWR). After the final models were generated, we added the Zn^2+^ back in all of them. We determined coordinates for the zinc atom by best-fit superimposing on Cα atoms the experimental Zn-bound structure with each CABS-flex model. For the TP53 DBD, we also used the pdb_mutate.py script from the HADDOCK utility scripts repository^90^ (https://github.com/haddocking/haddock-tools) to generate initial models for a set of protein variants from wild-type structure. We then applied CABS-flex to generate a structural ensemble for each variant, following the same procedure as for the wild-type protein.

### SNOmodel workflow and analysis of CABS-flex simulations

After obtaining the CABS-flex structural ensembles as detailed in the previous paragraph, we designed a Snakemake pipeline to analyze them called SNOmodels (https://github.com/ELELAB/SNO_investigation_pipelines). In particular, for each model of the previous step, we used MDAnalysis^86^ to calculate the χ_1_ dihedral angle of both SNO site and proximal cysteine residues, as well as the Cβ-Sγ-Sγ-Cβ between the two cysteine residues, similarly to what done for the TRAP1 crystal structures. We have also used PROPKA^85^ to predict the pKa of the Cysteine residues and used NACCESS ^83^ to calculate their relative solvent-accessible surface of side-chains. The pipeline provides both CSV output files and graphical representations of the results (deposited in OSF: https://osf.io/52bng/). For each protein target from SNOmodels we verified with AlphaFill^46^ that the SNO and proximal cysteines were not in proximity of a cofactor or metal ion. In the case of proximity to cofactor/metal binding sites, we discussed in detail in the result section. Indeed, the CABS-flex simulations have been performed without remodeling the cofactors because the coarse-grain approach does not support them. Indeed, according to the CABS-flex implementation, only protein groups are considered for the sampling. In the case of TP53, we remodeled the zinc ion in the binding pocket at the end of the run, and we applied structural restraints on the zinc-binding site. Similar approaches could be applied to other hits of interest in future studies, but we kept the simulation approach as simple as possible to analyze high-throughput hits.

### COSMIC analyses

We used the data provided in the available csv file of the Cancer Gene Census from COSMIC version 96 ^54,91^ to identify possible oncogenes and tumor suppressors for which we found at least a pair of an SNO site with a proximal cysteine with SNOfinder.

### MAVISp framework

We used the simple mode of the MAVISp framework^55^ and retrieved and aggregated the missense mutations for GRIN2A, CBFB, CALR, and EGFR in COSMIC ^54^, cBioPortal ^92^ and ClinVar ^57,58^. We verified if the selected target proteins contained intra-membrane regions by querying the Orientations of Proteins in Membranes (OPM) database^93^. In particular, we evaluated the position of SNO sites and proximal cysteines under investigation with respect to possible transmembrane regions. EGFR and GRIN2A were annotated in the OPM database with transmembrane regions predicted for the residues 646-721 (EGFR), along with the regions 556-582, 600-615, 618-648, and 812-839 (GRIN2A). Nevertheless, none of the cysteines of interest for our study are included in transmembrane regions. This step has been performed since the protocols for free energy calculations upon mutations applied in the MAVISp framework are currently not supporting transmembrane regions.

We used ClinVar Miner ^94^ to identify variants from ClinVar, whereas the aggregation step and the search for variants in COSMIC and cBioPortal have been carried out with Cancermuts ^95^. For TP53, we used Cancermuts to include also mutations from other sources ^56^. We then applied the STABILITY module of MAVISp to classify the effects of the mutations in unknown, destabilizing, stabilizing, or neutral according to changes in the folding free energy as estimated with the MutateX protocol ^96^ with Foldx5 ^97^, as well as with RosettaDDGPrediction^98^ with the cartddg2020 protocol ^99^ and the ref2015 energy function ^100^. We retrieved the classification for the variants of interest as benign, pathogenic, or variants of unknown significance from ClinVar ^58^. We used the classification provided by the STABILITY module of MAVISp to retain for further analyses only variants with unknown or neutral effects, as explained in the results ^55^.

### Analyses of CABS-flex simulations of mutant variants of TSG and OCGs

At first, we estimated with PyInteraph2^81,82^ the residues in the proximity of the SNO site and the proximal cysteine using the center-of-mass protein structure network approach and a distance cutoff of 8 Å. We then retained only those mutations from the MAVISp selection step affecting the residues in the proximity of the cysteine sites, which are the ones more interesting to explore for possible local effects on the *S*-nitrosylation mechanisms. The selection of the mutations to retain for analyses has been carried out with a Snakemake workflow, mutlist2SNO, available on GitHub (https://github.com/ELELAB/SNO_investigation_pipelines). Since PyInteraph2 applies distance criteria based on the center-of-mass of residue side chains, it does not consider glycines. For this reason, our workflow is not able to account for the effect of mutations on glycine residues in the proximity of the cysteines of interest. For the retained variants, we collected simulations with CABS-flex as described above, and we analyzed the structures using the pipeline SNOmodels described above.

### Plasmids

TRAP1 C261SC573R (C_2_) and C261SC501SC527AC57R (C_0_) were cloned in pET-26b(+) plasmid for bacterial expression using the Gene Synthesis & DNA Synthesis service from GeneScript Biotech (New Jersey, USA).

### Purification of recombinant TRAP1 variants

C_2_ and C_0_ TRAP1 recombinant proteins were produced in Origami (DE3) *E. coli* cells. Proteins production was induced with 1 mM [F1] IPTG 3h [F2] treatment (Isopropil-β-D-tiogalattopiranoside, VWR, Pennsylvania, USA). Then, proteins were purified using Ni-NTA resin (Quiagen, Germany, EU) according to manufacturer’s instructions. The protein release from the resin was achieved using a linear imidazole gradient generated by mixing buffer A (50 mM Phosphate Buffer, 250 mM NaCl, 10 mM imidazole, 10 mM β-mercaptoethanol, pH 7.8) and B (buffer A with 250 mM imidazole). Finally, protein-containing fractions were collected, combined, and dialyzed in the *storage buffer* (50 mM TRIS-HCl, 150 mM NaCl, 1 mm DTT, 1 mM EDTA, pH 7.5).

### In vitro treatments

Before treatments, DTT was removed from purified proteins by Zeba Spin Desalting Columns (Thermo Fisher Scientific, Massachusetts, USA), and proteins were transferred in *reaction buffer* (150 mM NaCl, 50 mM TRIS-HCl, pH 7.5). After protein quantification with DC Protein Assay Kit (Bio-Rad, California, USA), equal amounts of C_2_ and C_0_ TRAP1 were treated with or without 50 mM DTT (Sigma-Aldrich, Missouri, USA) to achieve total reduction, and 500 μM *S*-nitroso-*N*-acetylpenicillamine (SNAP) SNAP (Enzo Life Sciences, Switzerland), to induce *S*-nitrosylation. Reactions were performed at RT in the dark for 4 h.

### PEG-switch assay

PEG-switch assay was modified from previous studies ^101,102^. After treatment (performed as described above), excess DTT and SNAP were removed with Zeba Spin Desalting Columns and recombinant C_0_ and C_2_-TRAP1 mutants were transferred in *alkylation buffer* (100 mM TRIS-HCl pH 7.4, 1% SDS). Then, free thiols (in theory, only C501 and C527 of C_2_-TRAP1 variant) were blocked by incubating proteins with 10 mM *N*-ethylmaleimide (NEM, Sigma-Aldrich, Missouri, USA) for 30 min at 37°C in the dark. Excess NEM was, next, washed out with Zeba Spin Desalting Columns, and SNAP-modified thiols (to SNO or SS derivatives) were reduced by 50 mM DTT for 20 min at RT. After DTT removalwith Zeba Spin Desalting Columns, both C_0_ and C_2_-TRAP1 mutants were transferred in *Mal-PEG buffer* (100 mM TRIS-HCl pH 6.8, 0,5% SDS) and divided for labelling reaction. Labelling was performed with 4 mM maleimide-polyethylene glycol (Mal-PEG-5K) (Sigma-Aldrich, Missouri, USA) for 2 h, at RT, and stopped upon denaturing proteins with Laemmli SDS sample buffer (Thermo Fisher Scientific, Massachusetts, USA) and 12% β-mercaptoethanol (Sigma-Aldrich, Missouri, USA). Results were analyzed by Western blot using an anti-TRAP1 antibody (Santa Cruz Biotechnology, Texas, USA). Chemioluminescence was induced with Immobilon Western HRP Substrate (Merk-Millipore, Massachusetts, USA) and acquired with a digital camera (ChemiDoc, Bio-Rad, California, USA). Images adjustment (if any) was made with Fiji.

## Acknowledgements

The authors are grateful to Laila Fisher for her secretarial work.

EP group has been supported by Carlsberg Foundation Distinguished Fellowship (CF18-0314), Hartmanns Fond (R241-A33877), NovoNordisk Fonden Bioscience and Basic Biomedicine (NNF20OC0065262), and an HPC Grant DECI-PRACE 13th (CHAPREDO) on Archer (UK). MA is supported by a DDSA (Danish Data Science Academy) Ph.D. Fellowship (DDSA-PhD-2022-006). GF group has been supported by NovoNordisk Fonden Bioscience and Basic Biomedicine (2018-0052550), the Danish Cancer Society (KBVU, R146-A9414, and R231-A13855). EP and GF groups are part of the Center of Excellence in Autophagy, Recycling and Disease (CARD), funded by the Danish National Research Foundation (DNRF-125). CP was the recipient of a Ph.D. fellowship from the Danish Cancer Research Foundation (Dansk Kraeftforskningsfond, DKF-0-0-532). FF was supported by an AIRC “fellowship for Italy”.

## Author Contributions

Conception and Design: EP, GF; Development of computational methodologies: EP, MT; Development of experimental methodologies: AB, GF; Acquisition of data: EP, MT, ML, MA, MT, LC, KD, CP, FF, and FP. Analysis and interpretation of data: EP, MT, CP, GF. Writing of the manuscript: EP, MT, GF. Revision of the manuscript: all authors. Study supervision: EP, AB, GF.

