## Supplementary Figure S1 for "TRAP1 *S*-nitrosylation as a model of population-shift mechanism to study the effects of nitric oxide on redox-sensitive oncoproteins"

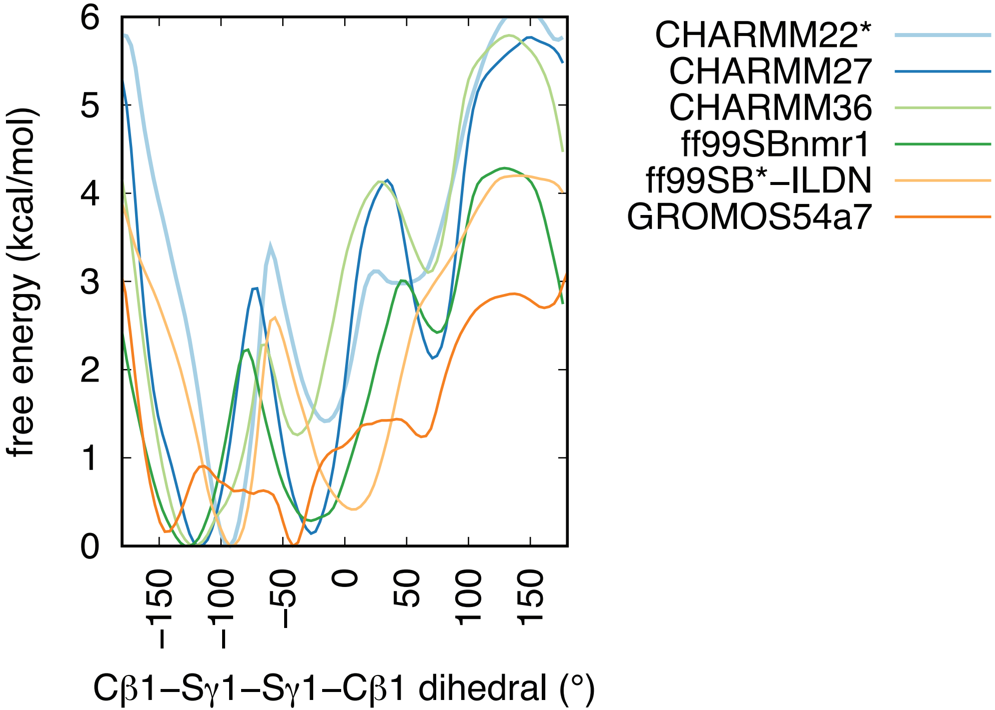


**Figure S1** Mono-dimensional free energy profiles for the Cβ-Sγ-Sγ-Cβ dihedral calculated on the metadynamics of *Danio rerio* TRAP1$_{311\text{-}567}$ with both SNO site and the proximal cysteine in their reduced form. We used eight different MD force fields. We observed that the different force fields overall captured the same main minima.
