## Supplementary Figure S2 for "TRAP1 *S*-nitrosylation as a model of population-shift mechanism to study the effects of nitric oxide on redox-sensitive oncoproteins"

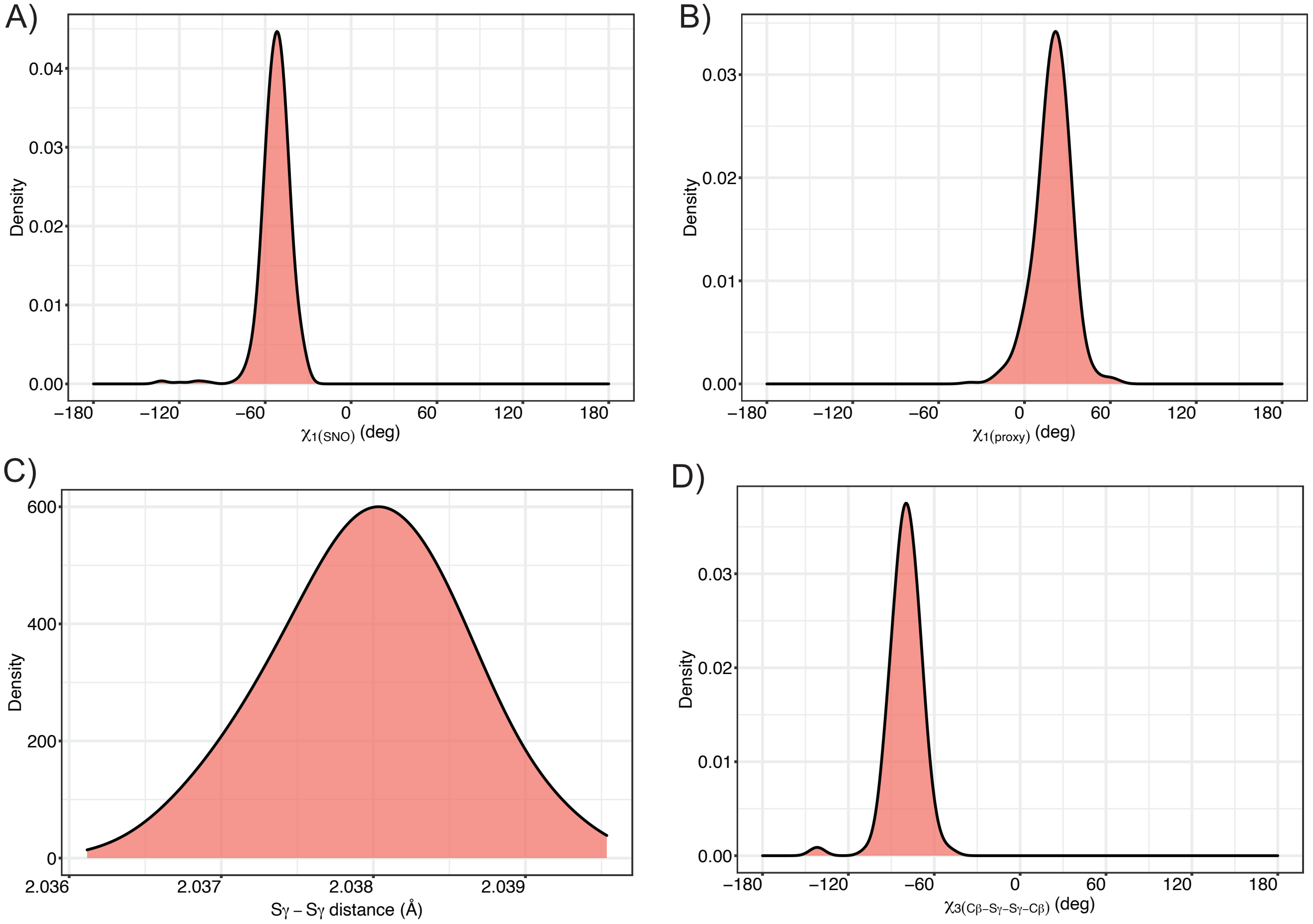


**Figure S2 Analysis of the unbiased MD simulation of TRAP1$_{227\text{-}438}$ with a disulfide bridge between C527 and C542.** We analyzed the collective variables used in metadynamics on a reference one-µs unbiased MD simulation of the oxidized form of TRAP1$_{311\text{-}567}$ with a disulfide bridge formed between C527, the S-nitrosylation site, and proximal C542. The plots show the distribution of the values calculated for the collective variables A) $\chi$1 dihedral of the S-nitrosylation site C527 ($\chi_{1(SNO)}$), B) $\chi$1 dihedral of the proximal C542 ($\chi_{1(proxy)}$), C) the distance between their sulfur atoms (Sγ-Sγ distance), D) the dihedral angle Cβ-Sγ-Sγ-Cβ. We observed that the Sγ-Sγ distance is approximately 2.03 Å. The Cβ-Sγ-Sγ-Cβ dihedral has a peak at -90 degrees which is common for disulfide bridges. In addition, we observed minus and plus states for the $\chi$1 of the S-nitrosylation and proximal cysteine, respectively.
