## Supplementary figures and images for "TRAP1 *S*-nitrosylation as a model of population-shift mechanism to study the effects of nitric oxide on redox-sensitive oncoproteins"

### Supplementary Figure S3

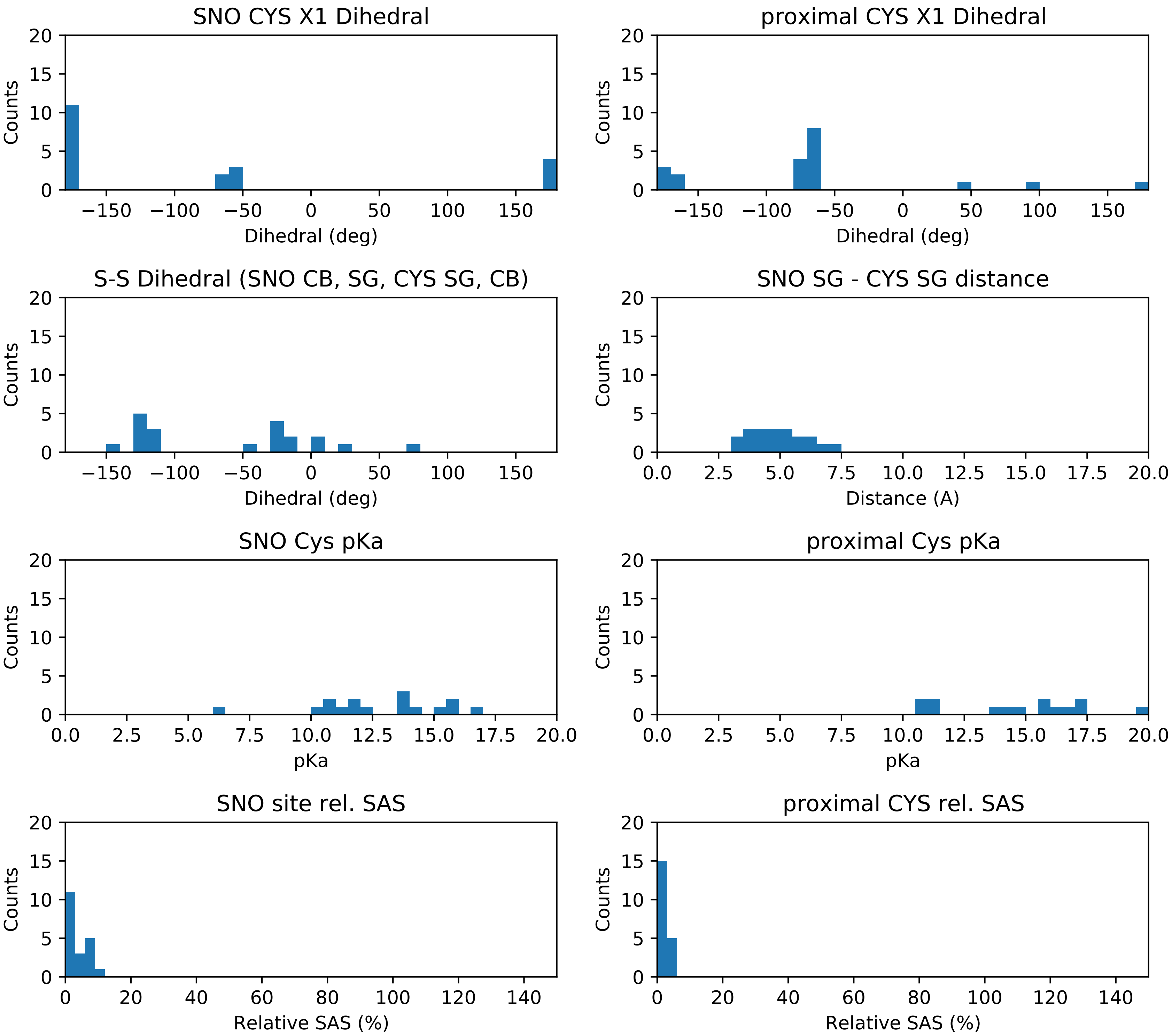
