## Supplementary Table S1 for "TRAP1 *S*-nitrosylation as a model of population-shift mechanism to study the effects of nitric oxide on redox-sensitive oncoproteins"

**Table S1. Overview of the structural parameters of the SNO site of TRAP1 and its proximal cysteines in structures from PDB.** The structures for the human and the *Danio rerio* variants are reported.

| **PDB entry** | **Cys ID (SNO site)** | **Cys ID (proximal)** | **Distance S-S**  (Å) | **Cβ-Sγ-Sγ-CβDihedral** (°) | **DSBASE2 Grade** | **χ_1 (SNO),_ χ_1(proxi)_**  (°) | **Resolution (Å)** |
| --- | --- | --- | --- | --- | --- | --- | --- |
| 4Z1G(A) | 501 | 527 | 2.6 | -118.8 | D | -65.0/55.0 | 3.10 |
| 4Z1H(A) | 501 | 527 | 2.1 | -124.4 | C | -160.9/-23.8 | 2.90 |
| 4Z1I(B) | 501 | 527 | 2.0 | -54.1 | C | -63.7/-25.0 | 3.30 |
| 4Z1I(C) | 501 | 527 | 2.0 | -86.2 | D | -62.7/46.7 | 3.30 |
| 4Z1I(D) | 501 | 527 | 2.0 | -66.0 | - | -62.1/47.2 | 3.30 |
| 5HPH(A) | 501 | 527 | 4.1 | -65.9 | - | -62.7/-35.1 | 2.43 |
| 5HPH(B) | 501 | 527 | 3.8 | -66.7 | - | -67.0/-24.9 | 2.43 |
| 5Y3N(A) | 501 | 527 | 5.1 | -149.0 | B | -173.3/-29.8 | 2.40 |
| 5Y3O(A) | 501 | 527 | 5.6 | -150.5 | B | 164.5/-31.5 | 2.70 |
| 4IPE(A) | 516 | 542 | 4.1; 2.0 | -54.5; -112.8 | - | -62.0/-40.9;-62.0/58.0 | 2.29 |
| 4IPE(B) | 516 | 542 | 4.2 | -61.0 | D | -60.5/-32.8 | 2.29 |
| 4IVG(A) | 516 | 542 | 4.2; 2.0 | -57.5; -112.1 | - | -65.9/-46.2;-65.9/67.4 | 1.75 |
| 4IYN(A) | 516 | 542 | 4.1; 2.0 | -56.7; -123.0 | - | -68.7/-44.5;/-68.7/55.6 | 2.31 |
| 4IYN(B) | 516 | 542 | 4.4; 2.1 | -64.7; -100.4; | - | -62.4/-32.7;-62.4/55.7 | 2.31 |
| 4J0B(A) | 516 | 542 | 2.0 | -94.1 | D | -63.0/45.1 | 2.35 |
| 4J0B(B) | 516 | 542 | 2.0;4.0 | -95.3;-65.5 | - | -61.4/53.7;-61.4/-21.0 | 2.35 |
| 5TTH(B) | 516 | 542 | 4.1 Å | -58.4 | D | -59.0/-32.0 | 3.20 |
| 5TVU(A) | 516 | 542 | 4.1 Å | -36.2 | B | -55.2/-51.4 | 3.50 |
| 5TVU(B) | 516 | 542 | 4.4 Å | -52.0 | D | -65.2/-48.1 | 3.50 |
| 5TVW(A) | 516 | 542 | 4.3 Å | -52.0 | D | -62.5/-44.0 | 2.50 |
| 5TVW(B) | 516 | 542 | 4.4 Å | -55.3 | D | -65.4/-42.1 | 2.50 |
| 5TVX(A) | 516 | 542 | 4.3 Å | -50.9 | D | -63.5/-49.7 | 2.20 |
| 5TVX(B) | 516 | 542 | 4.5 Å | -62.2 | - | -66.5/-35.2 | 2.20 |
| 6D14(A) | 516 | 542 | 4.0 Å | -55.2 | - | -58.1/-34.3 | 2.50 |
| 6D14(B) | 516 | 542 | 4.0 Å | -54.2 | D | -61.2/-27.6 | 2.50 |
